# A critical look at directional random walk modeling of sparse fossil data

**DOI:** 10.1101/2025.11.11.687792

**Authors:** Rolf Ergon

## Abstract

The general random walk model (GRW) of Hunt (2006) is used to infer directional evolution in mean trait values from sparse fossil data. Such evolutions are modeled as the accumulated result of small steps with mean step sizes and step variances. As shown in simulations and real data cases, the mean step sizes are often easy to estimate from data, except for cases where the mean step size is small compared to the step variance. The step variances, on the other hand, can be estimated reasonably well only when the mean trait values have small measurement errors, but even here the step variance estimation may be difficult. For fossil data with realistic measurement errors, the step variances appear to be extremely difficult to find, and they are often found to be negative. They must then be set to zero, such that GRW collapses into deterministic walk processes plus sampling errors. As a result of poor step variance estimates, the directional evolution may be both under- and overestimated as compared with generalized least squares (GLS) results, which gives the best linear unbiased estimator (BLUE) of the evolutionary slope. Here, I study this problem through simulations as well as in four real data cases. Based on weighted mean square error (WMSE) comparisons, my conclusion is that GLS in cases with large measurement errors is the best method for inference of directional evolution. When the step variances must be set to zero, GLS is simplified into weighted least squares (WLS) estimation.

## 1 Introduction

Hunt (2006) developed what he called a general random walk (GRW) model for analyses of fossil time series, later referred to as a model for directional evolution (Hunt, 2012). This model assumes a random walk process where at each timestep an increment of evolutionary change in a mean trait value is drawn at random from a distribution of evolutionary steps, and that this incremental change is normally distributed with mean value *µ*_*step*_ and variance 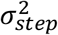. For an observed evolutionary mean trait change Δ*Y* over T time steps the log-likelihood function for a GRW process is given by (Hunt, 2006)

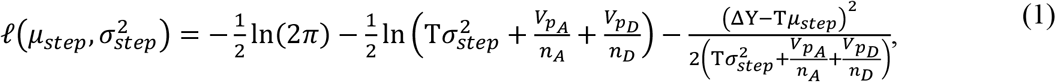

where *n*_*A*_ and *n*_*D*_ are the numbers of observed ancestors and descendants, respectively, while 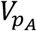 and 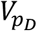 are the corresponding population phenotypic variances. With *N* irregular and sparse samples of mean trait values, multiple ancestor-descendant trait differences over *N* – 1 evolutionary transitions may be used jointly to estimate *µ*_*step*_ and 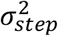 by summing the log-likelihoods according to Eq. (1) over the transitions, i.e., by use of 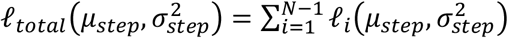. From this expression step parameters can be found by means of maximum likelihood estimation, and the estimated step size 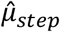 will then be a measure of directional change over time.

For a single transition it is obvious from Eq. (1) that 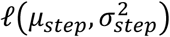 is maximized when 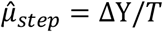, and as shown in simulations in Section 3, *µ*_*step*_ can be well estimated also from data over several transitions. From Eq. (1) it is also obvious that it is difficult to estimate 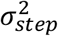 in cases where 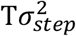 is small compared to *V*_*p*_/*n* for the samples.

When *µ*_*step*_ is fixed to zero we obtain an unbiased random walk (URW) process, and the estimated step variance 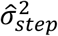 then becomes a scaled version of the rate of directional change (Hunt, 2012). I will not make use of this property, but it should be clear that the estimation difficulties will not disappear when *µ*_*step*_ is set to zero.

Hunt (2006) presented simulation results over 19 transitions with clearly directional evolution, where both *µ*_*step*_ and 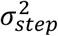 were estimated well, except for small values of 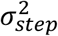 as compared with 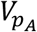 and 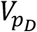. The population phenotypic variances in these simulations appear to be unrealistically small, as compared with real data cases in Section 4. In Hunt (2012) similar small values appear in Figure 2, panel A, illustrating directional evolution. In simulations in Section 3 I essentially repeat parts of Hunt’s simulations with what appears to be more realistic phenotypic variances, and in around 40% of the realizations I find that 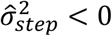, which in practice means that 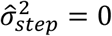. In such cases the GRW model collapses into a deterministic directional walk model plus sampling errors.

Eq. (1) is not primarily intended for predictions, but once 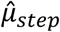 has been determined a prediction model 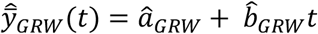 is found by use of 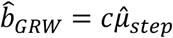, and by fitting *â*_*GRW*_ to the data by a generalized least squares approach. Here, *c* is a scaling constant that can be chosen freely (Hunt, 2006), and throughout the paper I will set *c* = 1.

In my simulations I also analyze data for directional evolution by means of the generalized least squares (GLS) method, which gives the best linear unbiased estimator (BLUE) of the evolutionary slope. I then find that GRW may both underestimate and overestimate prediction slopes by up to 50%. For comparisons of GRW and GLS slope estimates, the simulation results include weighted mean square errors (WMSE).

In Section 4 I analyze data from four real data cases, a bryozoan case, two ostracod cases, and a stickleback fish case. In all these cases parameter estimation with 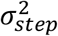 as a free variable resulted in 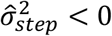, which means that 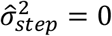 must be used in a realistic model, and as a result of this GLS is simplified into weighted least squares (WLS).

In the first three cases there exists a dominating evolutionary driver, and a tracking model as discussed in Ergon (2025a) is then an attractive alternative. The applicability of such a model will, however, depend on the quality of the evolutionary data, and only in the bryozoan case is the tracking model better than the WLS model. As shown in Ergon (2025b) it is also in the stickleback fish case possible to find a tracking model that is clearly better than the WLS model, but then by use of an association with rather than a direct measurement of an environmental proxy. Ergon (2025b) also includes improved tracking models for the two ostracod cases, using a different temperature proxy.

My results are summarized and discussed in Section 5, and my conclusion is that GLS or WLS are the best alternatives for analyses of directional evolution, and that dependent on the quality of the data a tracking model may give even better prediction results. Appendix A gives some details regarding the tracking model for the stickleback fish case. Some detailed simulation results are given in Appendix B, results for various values of 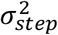 are given in Appendix C, while Appendix D gives data for the four real data cases.

## 2 Methods

### 2.1 GRW predictions

Eq. (1) is not intended for predictions, but once 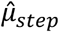 is determined a prediction model

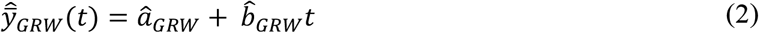

is found by use of 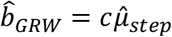, and by fitting a parameter *a*_*GWR*_ to the data by a generalized least squares approach. Here, *c* is a scaling constant that I consistently will set to *c* = 1.

### 2.2 Methods for simulations

In the simulations in Section 3 I first repeat some of the simulations in Hunt (2006), but with the number of samples reduced from 20 to 10. I generate time series data over 1,000 time steps as 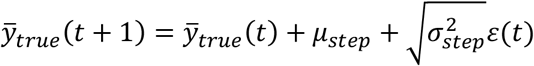, where *ε*(*t*) is a normally distributed random number with variance one, and I then extract ten samples, 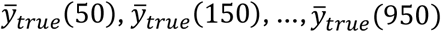. The initial value is chosen as 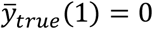. I use *µ*_*step*_ = 0.1 and 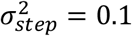, which over 1,000 timesteps results in an approximately linear change of 100 in 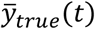. To each of the ten samples I add the mean values of *n* = 30 individual measurement errors drawn from normally distributed populations with mean *µ*_*p*_ = 0 and variance *V*_*p*_ = 1, and thus form ten samples 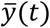. Each of these samples has a standard error 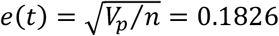, i.e., a small error compared to the total change in 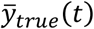 and thus in 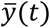.

Based on the generated 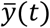 data with equal *V*_*p*_/*n* values for all samples, I find estimates 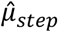 and 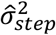 that maximize the log-likelihood function 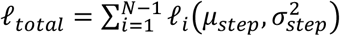, where *l*_*i*_ is given by Eq. (1). For this purpose, I use the *fmincon* function in MATLAB, i.e., I minimize 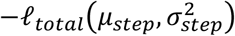.

In order to obtain standard errors comparable to real data, as presented in Section 4, I repeated the simulations with *V*_*p*_ = 400. In addition I also let the sampling times be irregular as uniformly distributed random numbers in the ranges 1 − 100, 101 − 200, …, 901 − 1000, while the numbers of individual samples varied between 3 and 60 with uniform random distributions.

### 2.3 Covariance matrix for random walk process

In order to find the covariance matrix ***C*** of a random walk process we may for simplicity assume *µ*_*step*_ = 0, such that the expected value at all time steps is zero. For an element *c*_*t,s*_ in ***C*** with *s* > *t* we will then find

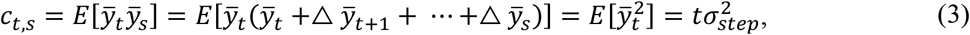

where *E*[·] is the expectation operator, and where the incremental changes 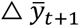 etc. are independent of 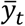. The result from Eq. (3) is the well-known variance of integrated discrete white noise. With *N* samples we thus find

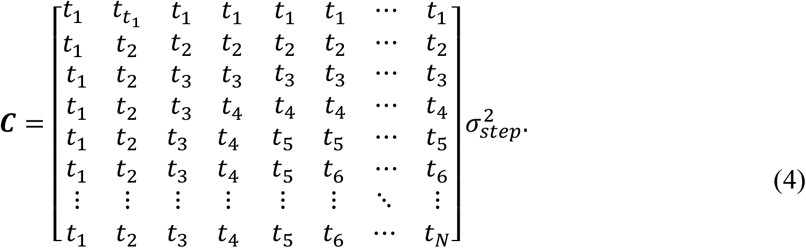

With *µ*_*step*_ ≠ 0 we will have a combination of directional evolution and integrated white noise, which is a clear parallel to the combination of natural selection and genetic drift as discussed in Walsh and Lynch (2018), Ch. 18.

### 2.4 Generalized and weighted least squares estimation

In the simulations, predictions 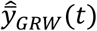 according to Eq. (2) will be compared with generalized least squares (GLS) predictions found as 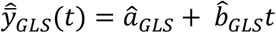. Here, *â*_*GLS*_ and 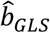 are found as

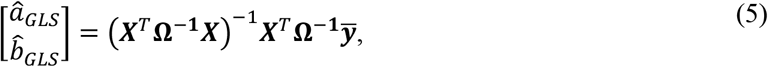

where 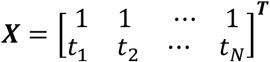 and 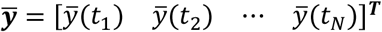, while **Ω** = ***C*** + ***V***, where ***V*** is a diagonal matrix with elements *v*_*t*_ = *V*_*p*_/*n*_*t*_. Note that the prediction slope parameter 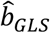 is the best linear unbiased estimator (BLUE).

Comparisons of responses 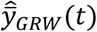 and 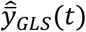 can be done by use of generalized mean squared errors

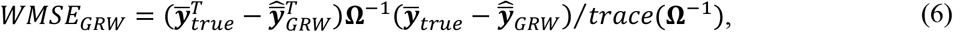

and

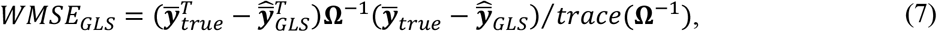

where 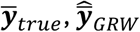 and 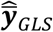 are *N* × 1 data vectors. Note that the division by *trace*(**Ω**^−**1**^) makes the WMSE results consistent with MSE for ordinary least squares (OLS), where **Ω** = ***I*** and *trace*(**Ω**^−**1**^) = *N*, but this normalization is of no importance for comparisons of GRW and GLS results. Also note that in real data cases 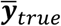 must be replaced by the observation vector 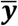.

In all four real data cases in Section 4 the realistic step variances are found to be zero, such that ***C*** = 0. From Eq. (5) then follows the weighted least squares (WLS) result

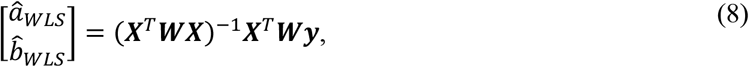

where ***W*** = ***V***^−**1**^. From this follows that **Ω**^−1^ in Eqs. (6) and (7) is replaced by ***W***, while *trace*(**Ω**^−1^) becomes 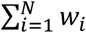.

## 3. Simulation results

As in Hunt (2006), GRW parameter estimation results with phenotypic variance *V*_*p*_ = 1 were found from 1,000 repeated simulations, with typical responses from a system with *µ*_*step*_ = 0.1 and 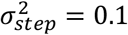 as shown in Fig. 1, left panel. In the same simulations, GRW and GLS prediction parameters *â*_*GRW*_ and 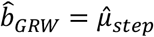 as well as *â*_*GLS*_ and 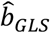 were found, which gave typical responses as also shown in Fig. 1, left panel. The results are summarized in Table 1, including *WMSE* values based on normalized weights. The 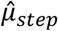 and 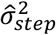 results are very much the same as found by Hunt (2006), and also here 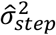 consistently approaches 0.1(*N* − 1)/*N* = 0.09. Note that the measurement standard errors as shown by error bars are quite small. Also note, however, that the SE value for 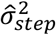 is 50% of the mean value, indicating the estimation difficulty also in the simple case with *V*_*p*_ = 1.

**Table 1.**
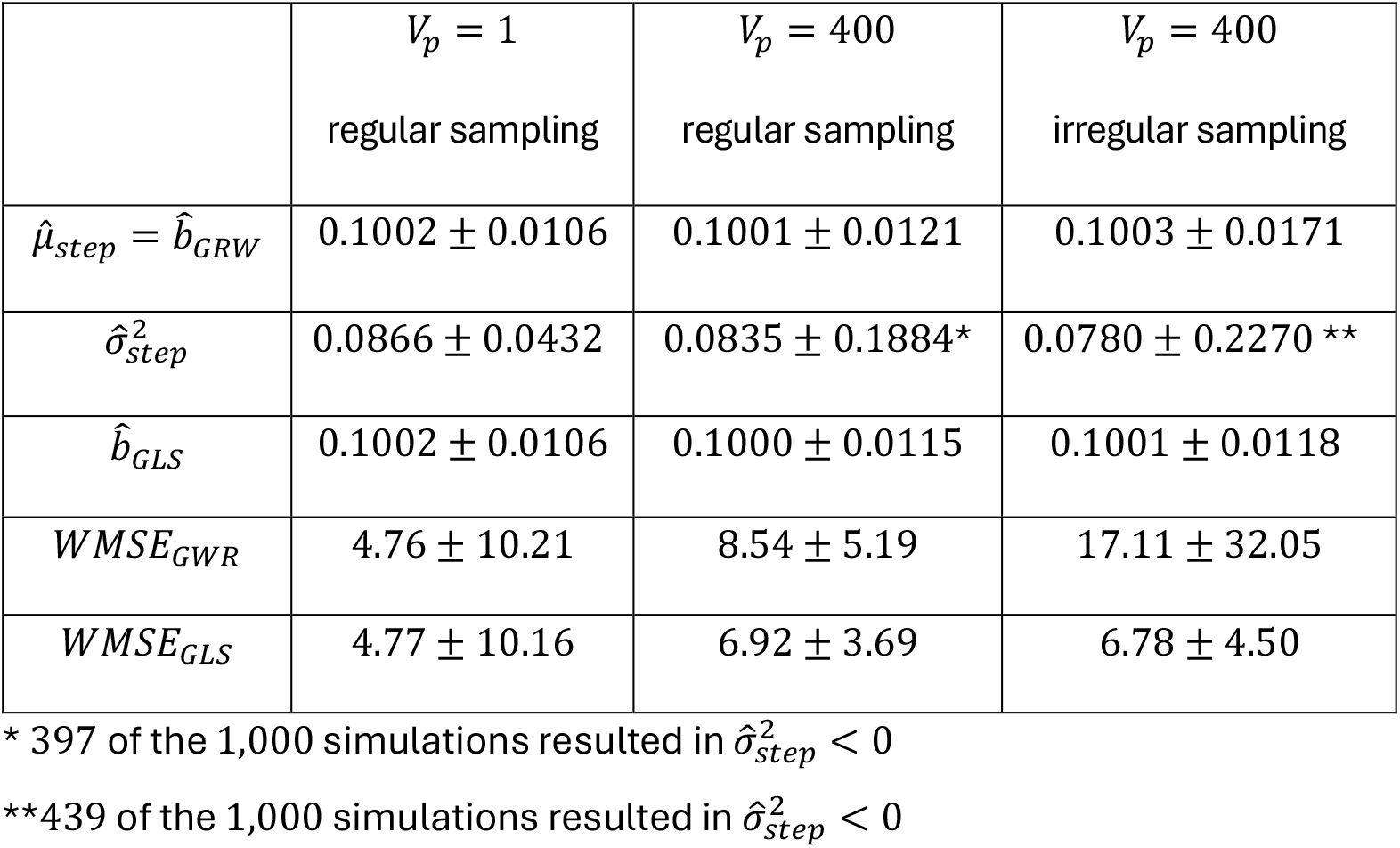
Simulation results given as *mean* ± *SE* from *1,000* realizations.

**Figure 1.**
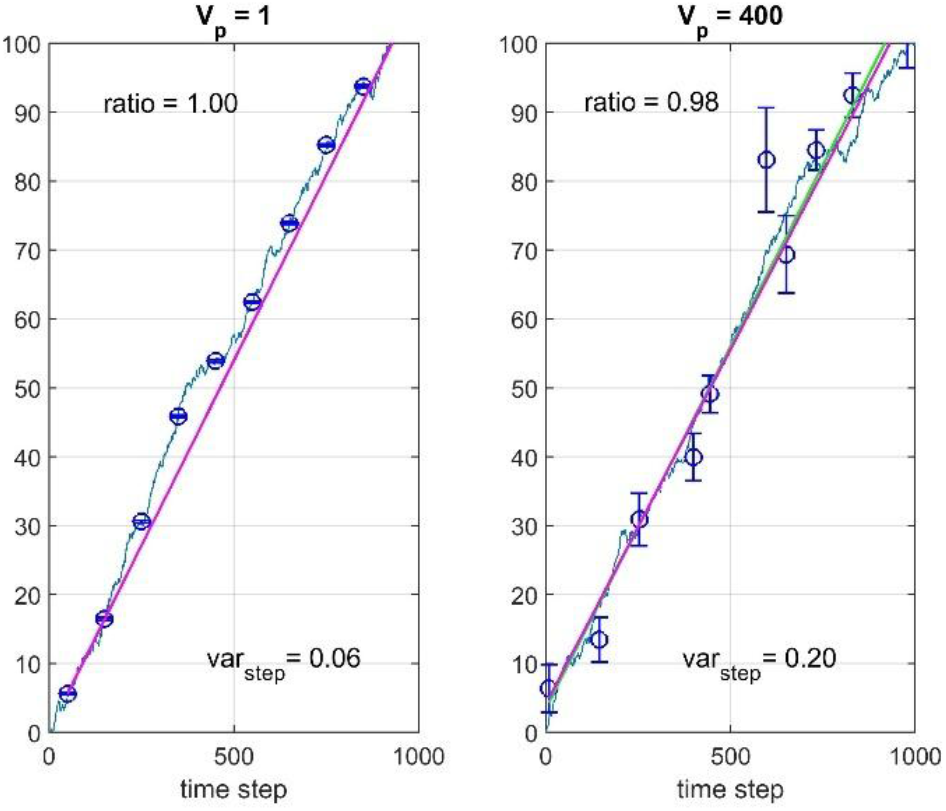
Typical results for simulations with *µ*_*step*_ = 0.1, 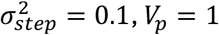 and regular sampling (left panel) and with *V*_*p*_ = 400 and irregular sampling (right panel). Values of 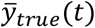 are shown by solid blue lines, while 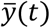 are shown by circles with error bars. GLS predictions are shown by green lines, and GRW predictions are shown by red lines. Estimated step variances 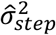 and slope ratios 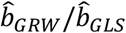 are shown in the plots.

**Figure 2.**
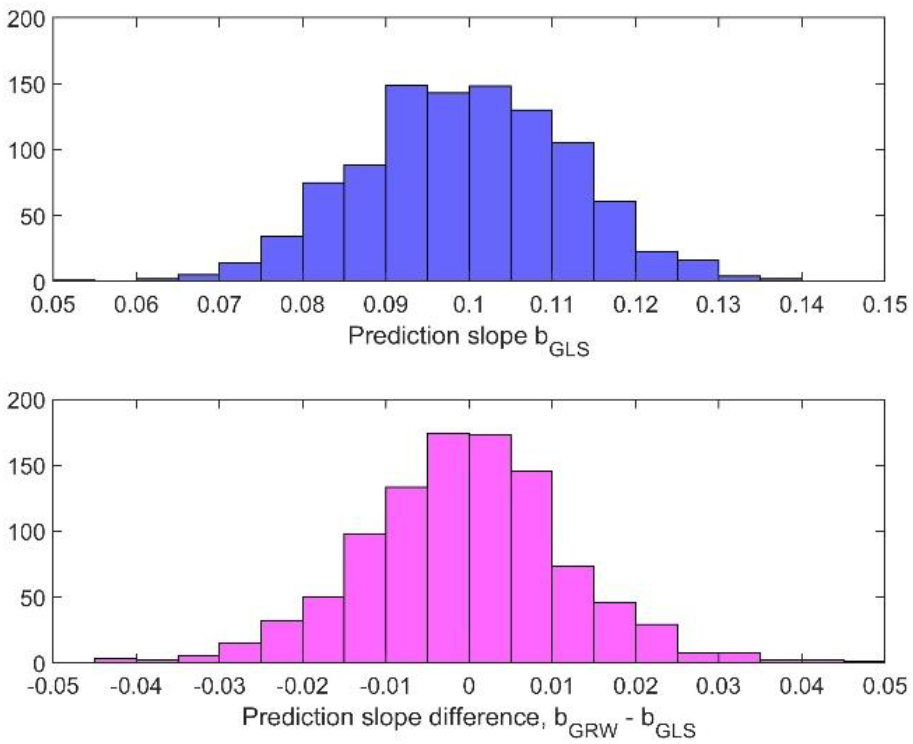
Typical histograms for estimated prediction slope 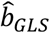 (upper panel) and for the GRW estimation error 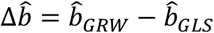 (lower panel), based on 1,000 realizations with *µ*_*step*_ = 0.1, 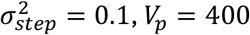, irregular sampling and variable numbers of samples as described in text.

In order to obtain standard errors comparable to the ones in the real data examples in Section 4, we must increase *V*_*p*_ considerably. Simulation results with *V*_*p*_ = 400, regular sampling, and equal numbers of individuals for all measurements are also shown in Table 1. Note that the SE value for 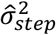 now is much larger than the mean value, and that 397 of the 1,000 realizations resulted in 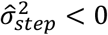. In such realizations, *WMSE*_*GWR*_ was computed by use of 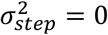. Results with irregular sampling and variable numbers of individual samples, as described in Section 2, are included in Table 1, and typical responses are shown in Fig. 1, right panel. Note that the estimates of *µ*_*step*_ are very much the same when *V*_*p*_ is increased from 1 to 400, but that 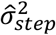 is very much affected. More detailed parameter estimation results are given in Appendix B.

The *WMSE* values are as seen in Table 1 increased when *V*_*p*_ is increased from 1 to 400, but the essential differences from the simulations in Hunt (2006) are still the poor estimation results for the step variances. As a result, the slope of directional evolution as found from 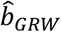 can be 50% under- or overestimated, as compared with 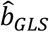, see histogram in Fig. 2 (lower panel). Note that the GRW estimation error 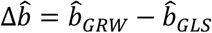 with 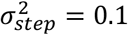 has approximately the same variance as the estimated slope 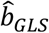 (upper panel). As shown in Appendix C, the variance of 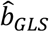 increases with 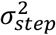, while the variance of the GRW estimation error decreases. Three typical realizations are shown in Fig. 3.

**Figure 3.**
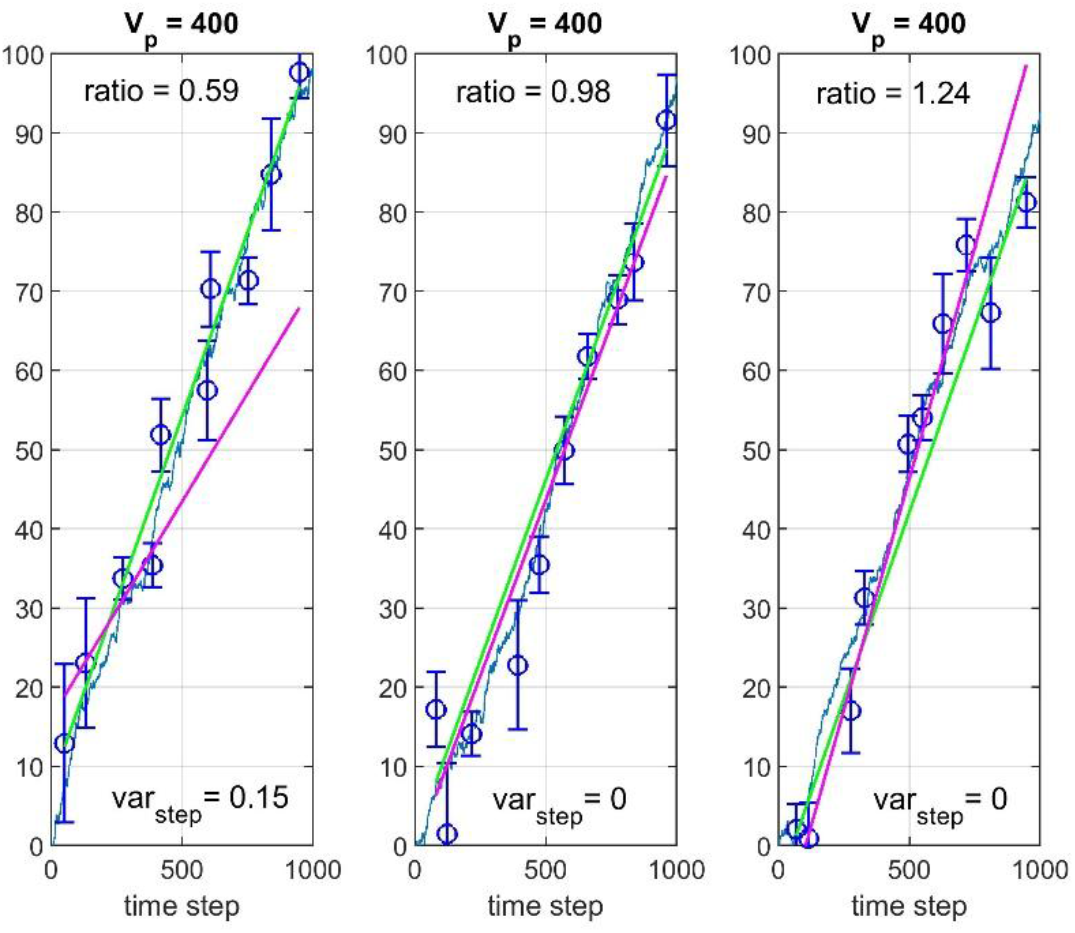
Prediction results for three realizations with *µ*_*step*_ = 0.1, 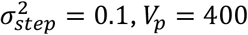, irregular sampling and variable numbers of samples as described in text. Values of 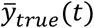 are shown by solid blue lines, while 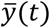 are shown by circles with error bars. GLS and GRW predictions are shown by green and red lines, respectively. Estimated step variances 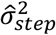 and slope ratios 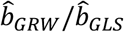 are shown in the plots.

As an illustration of the difficulties involved, Appendix C also shows responses for two quite different evolutionary realizations of a system with *µ*_*step*_ = 0.1 and 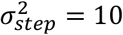, both apparently directional, but with different signs. Also in these cases, however, *WMSE*_*GLS*_ is clearly lower than *WMSE*_*GRW*_.

## 4 Real data cases

### 4.1 Bryozoan case

Liow et al. (2024) presented results from a field study of the bryozoan species *Microporella agonistes*, based on a fossil record over 2.3 million years. Here, I will use the time series for *log AZ area mean* as shown in Fig. S2, panel A, in Liow et al. (2024), with the two last data points omitted (because they do not fit into a linear model, see Ergon (2025a) for details). In this case there are no individual data points, but instead colony means, and the mean trait values are thus means of colony mean values.

As shown in Ergon (2025a), good predictions can be found using an adaptive peak tracking model based on a moving average smoothed version of the ∂^18^*O*(*t*) environmental proxy, with window size 100. An essential feature of this model is that the mean trait values as function of ∂^18^*O*_*smooth*_ can be approximated by a straight line, see Fig. 9 in Ergon (2025a). When this line is found by WLS the result is a tracking model with known WMSE values. The ∂^18^*O*(*t*) data are found in Lisiecki and Raymo (2005), while other data are collected in Table D1 in Appendix D.

Since GWR without parameter limitations resulted in 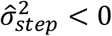, I used 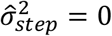, such that GLS is simplified into WLS. Fig. 4 shows predictions by means of the tracking model as well as WLS and GRW predictions. Here, the prediction slope 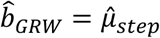, i.e., I use 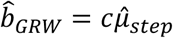 with *c* = 1. Prediction slope and WMSE results for the three models are given in Table 2. Note that the WLS results are better than the GRW results, but that the tracking results are even better.

**Table 2.** Results for real data cases.

|  | Bryozoan | Ostracod 1 | Ostracod 2 | Stickleback |
| --- | --- | --- | --- | --- |
| $\hat{b}_{WLS}$ | 0.1519 | 0.0402 | 0.0434 | 1.2605 |
| $\hat{b}_{GRW}$ | 0.1441 | 0.0316 | 0.0255 | 1.2531 |
| $WMSE_{Tracking}$ | 0.00100 | 0.000279 | 0.000613 | — |
| $WMSE_{WLS}$ | 0.00190 | 0.000241 | 0.000606 | 0.00040126 |
| $WMSE_{GRW}$ | 0.00193 | 0.000319 | 0.0011 | 0.00040132 |

**Figure 4.**
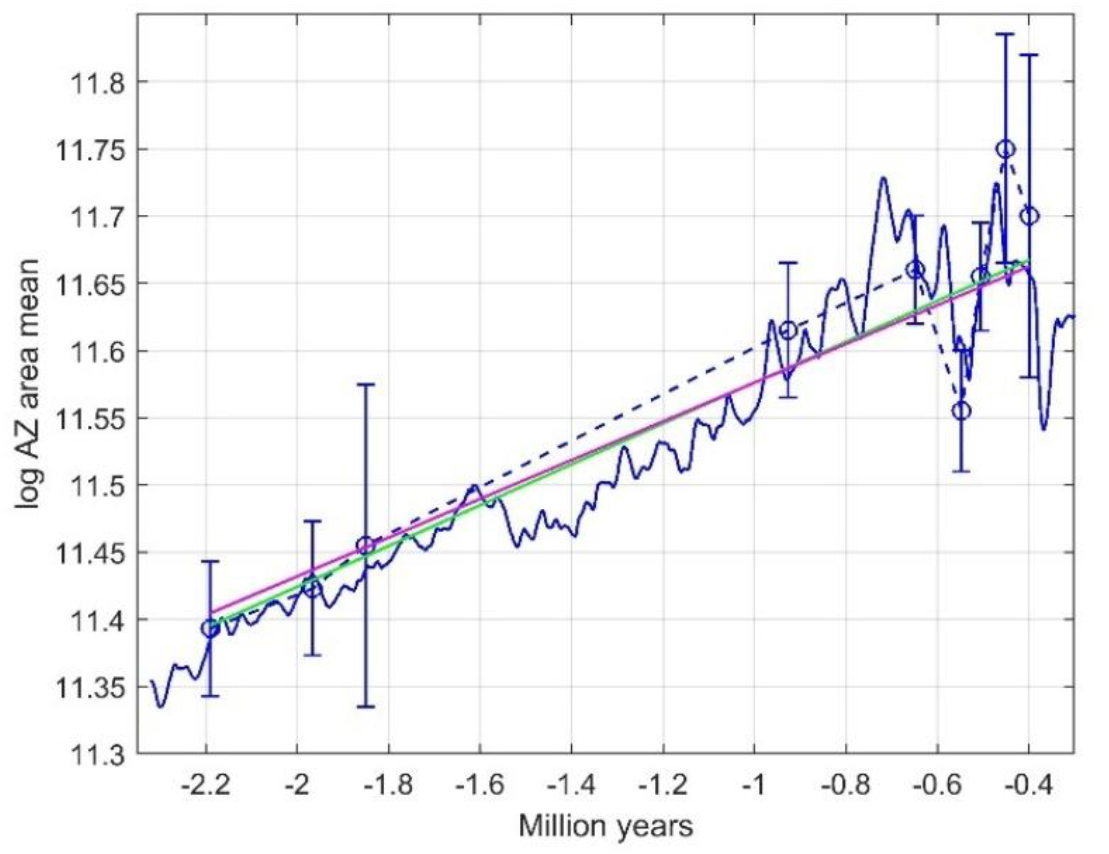
Tracking, WLS and GRW results for Bryozoan case, with mean trait observations (dashed blue line with circles and error bars), predictions based on a centered moving average smoothed version of the ∂^18^*O*(*t*) temperature proxy (solid blue line), WLS predictions (green line), and GRW predictions with 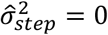 (red line).

#### 2 Ostracod case 1

Hunt and Roy (2006) presented time series data for body size of the ostracod genus *Poseidonamicus*, and in this case I use data for *P. major*, extracted as shown in Table D2 in Appendix D. The data include long-term smoothed Ma/Ca temperature data and the number of samples in each population. Mean valve length is measured in *µm*, with measurement variance estimated as 467 (*µm*)^2^ (Hunt et al., 2015). In the data analysis the trait values are log-transformed, with the measurement errors transformed accordingly, i.e., 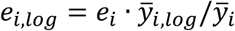. The original and log-transformed samples thus have equal coefficients of variation.

As in the Bryozoan case, the mean trait as function of the temperature proxy could be approximated by a straight line, such that a tracking model could be used. Also in this case the realistic step variance is found to be 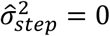, such that GLS is simplified into WLS.

Fig. 5 shows predictions by means of the Mg/Ca tracking model as well as of WLS and GRW models. Prediction slope and WMSE results are given in Table 2. Note that the WLS model is better than the tracking model, presumably owing to age uncertainty in the Mg/Ca data, and that the GRW model underestimates the prediction slope by 21%. Verification of the search results are shown in the grid contour plot in Fig. 6, where it should be noted that 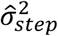 is allowed to be negative.

**Figure 5.**
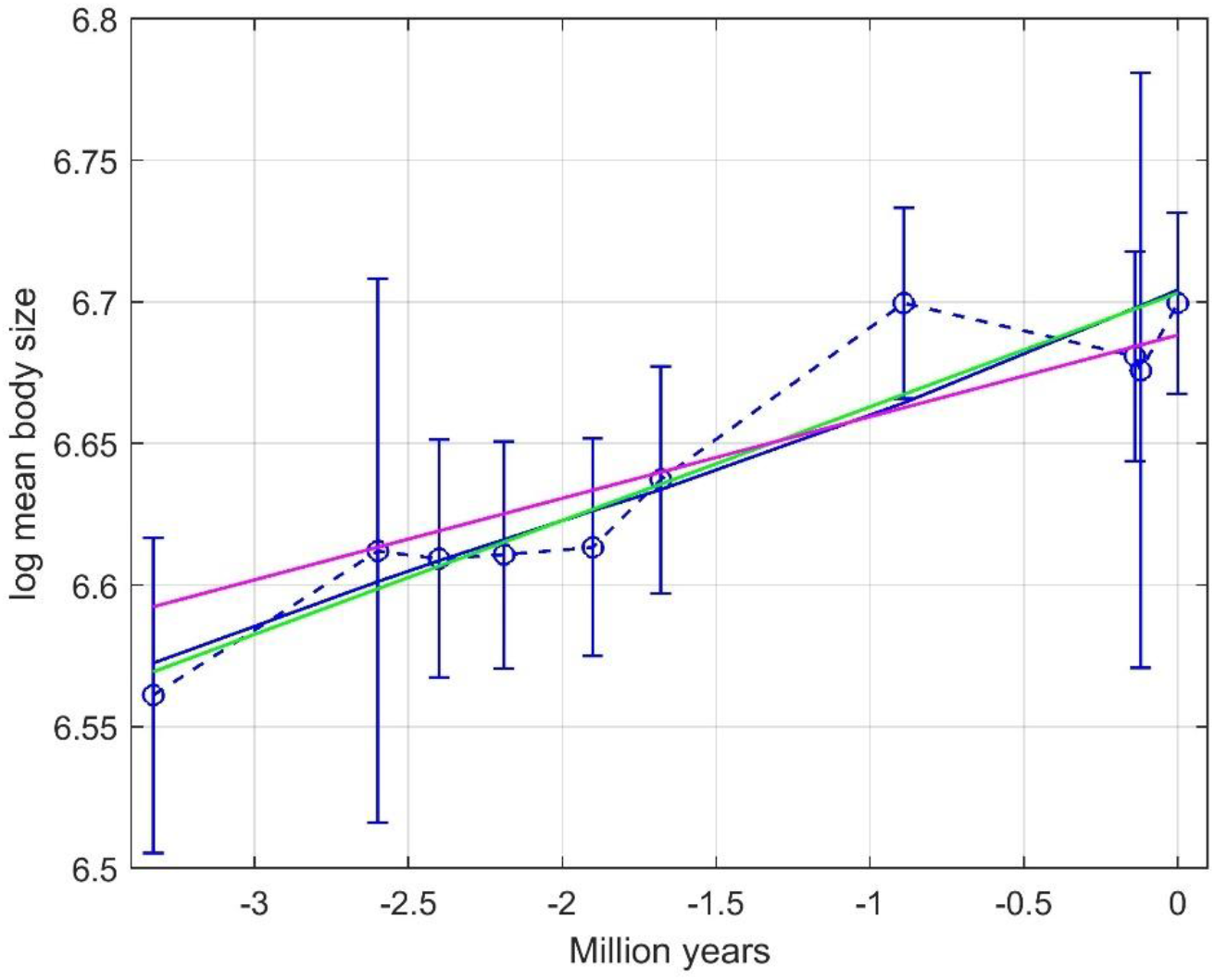
Results for Ostracod case 1, with mean trait observations (dashed blue line with circles and error bars), predictions based on the Mg/Ca temperature proxy (solid blue line that is not quite a straight line), WLS predictions (green line, almost identical to the solid blue line), and GRW predictions with 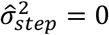 (red line).

**Figure 6.**
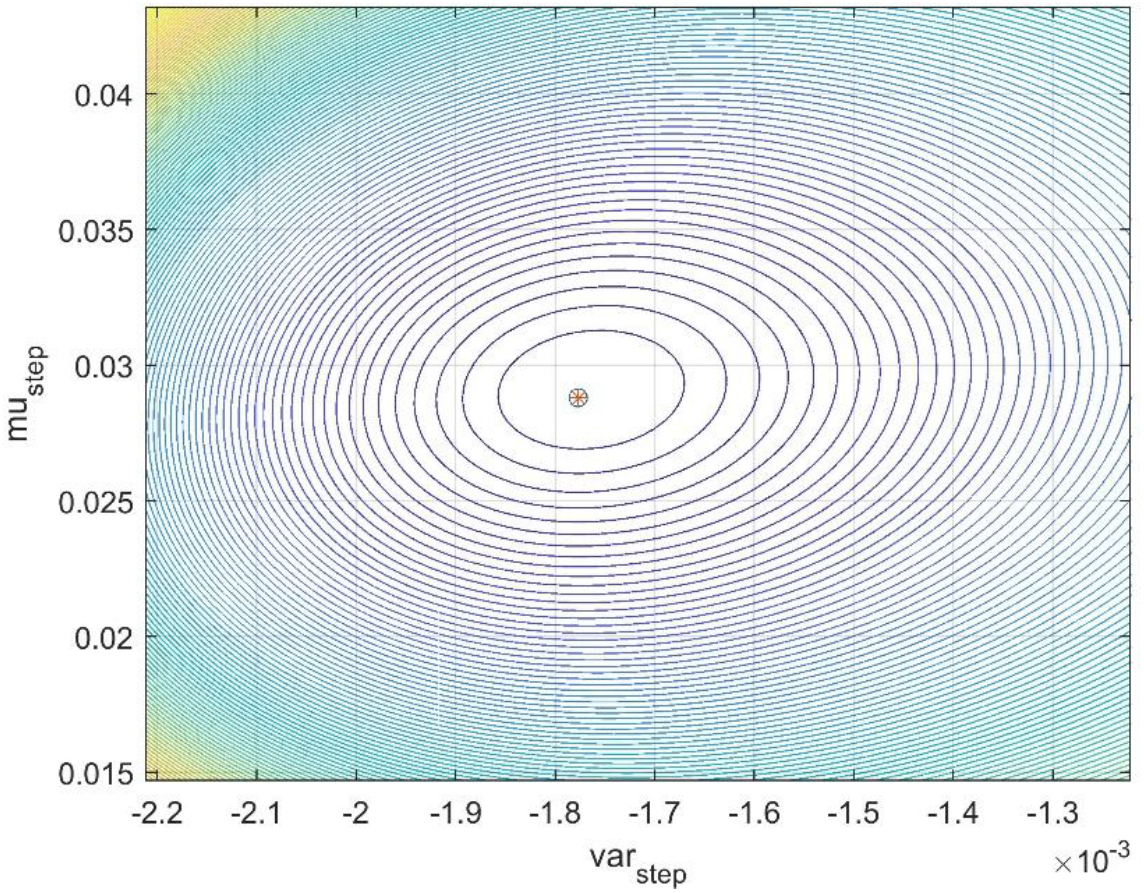
Grid contour results for Ostracod case 1, with the point 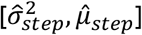 added as circle with star. Note that 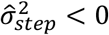 is accepted.

### 4.3 Ostracod case 2

In this case I use data for *P. pintoi* in Hunt and Roy (2006), as shown in Table D3 in Appendix D. This is another case where the realistic step variance is found to be 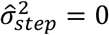. As in the Ostracod case 1, the mean trait as function of the temperature proxy could be approximated by a straight line, such that a tracking model could be used.

Fig. 7 shows predictions by means of a Mg/Ca tracking model as well as of the WLS and GRW models. Prediction slope and WMSE results are given in Table 2. Note that also here the WLS model is better than the tracking model, presumably owing to age uncertainty in the Mg/Ca data, and that the GRW model underestimates the prediction slope by 41%. As a verification of the search results, Fig. 8 shows the log likelihood as function of *µ*_*step*_ with 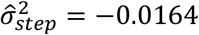 (the result with 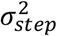 as free variable, upper panel) and 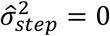 (lower panel). As seen in Fig. 8, the maximum log likelihood value is reduced when the search is limited to positive values of 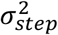, i.e., when 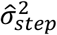 is non-optimal.

**Figure 7.**
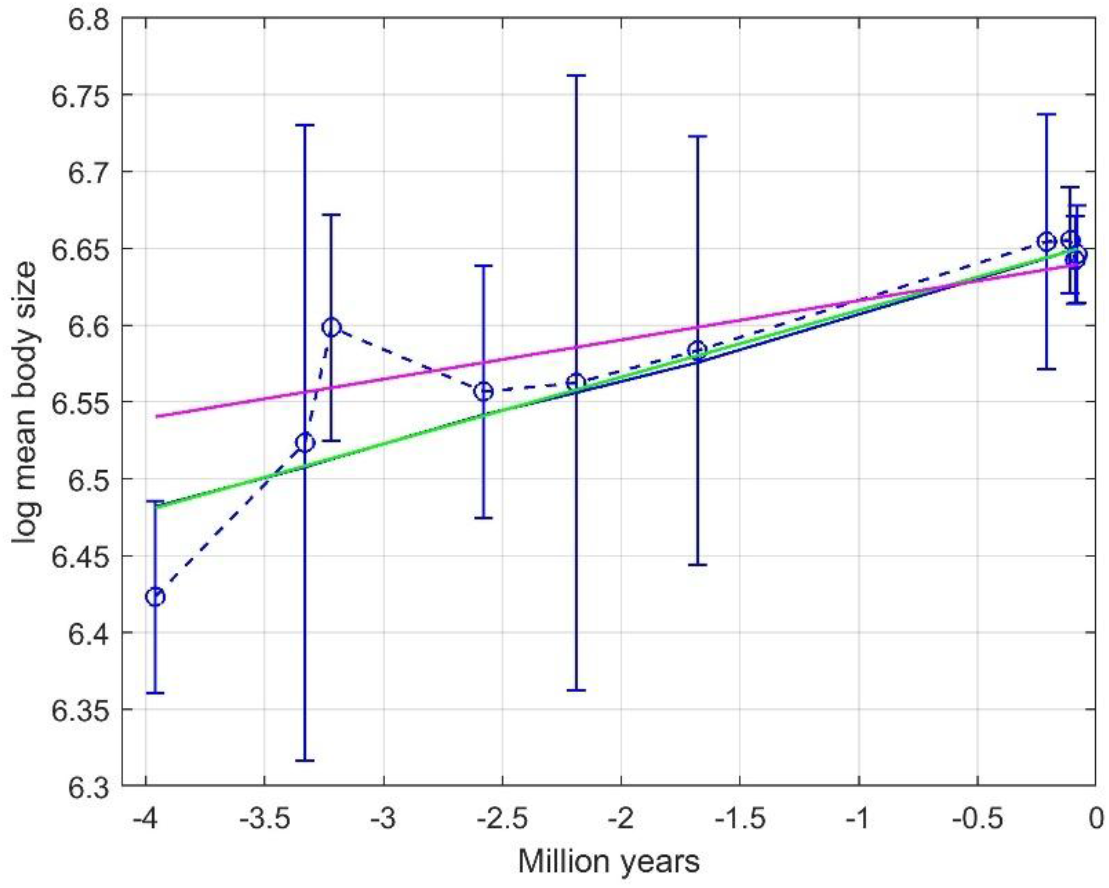
Results for Ostracod case 2, with mean trait observations (dashed blue line with error bars), predictions based on the Mg/Ca temperature proxy (solid blue line that is not quite a straight line), WLS predictions (green line, almost identical to the solid blue line), and GRW predictions with 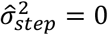 (red line).

**Figure 8.**
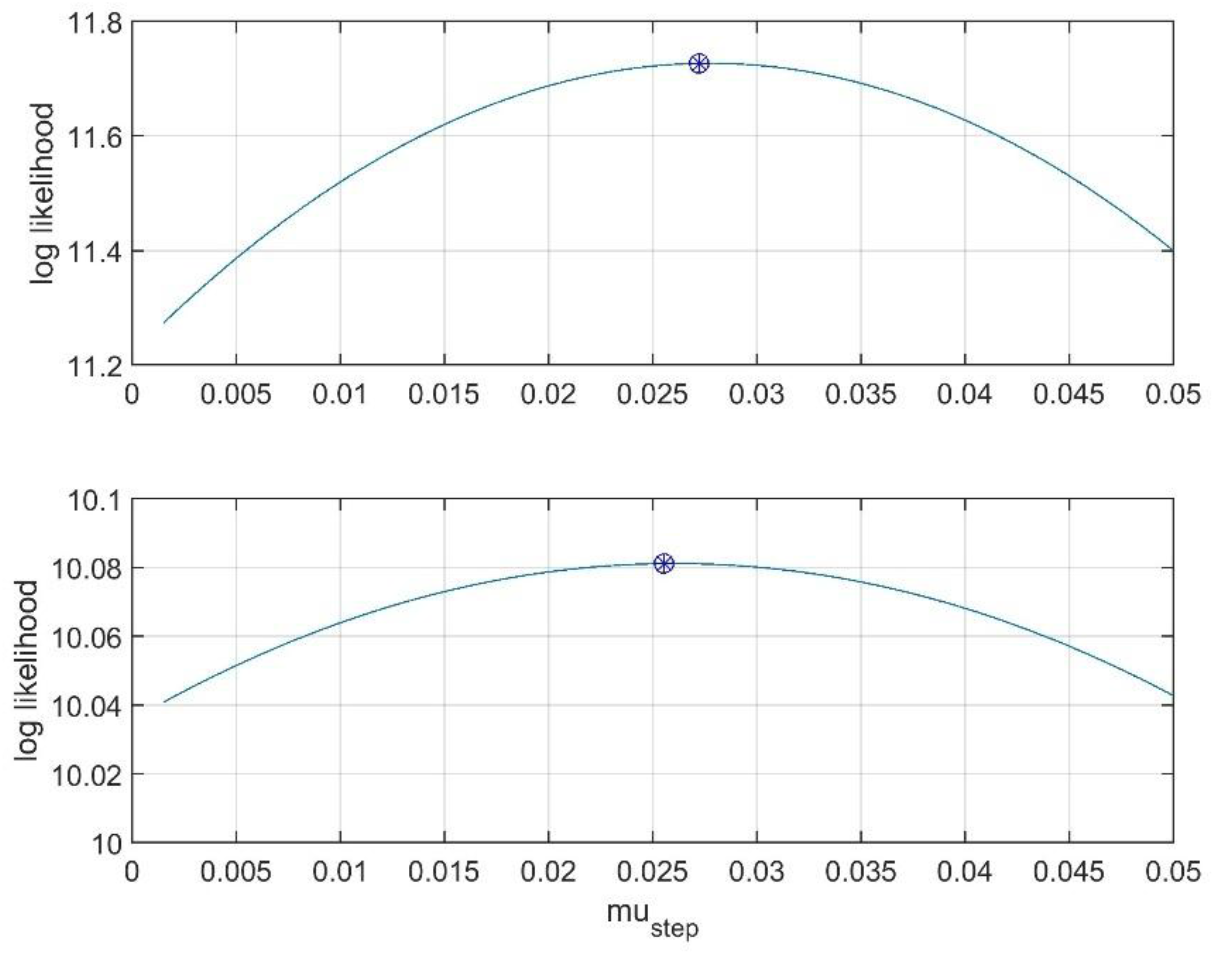
The log likelihood as function of *µ*_*step*_ with 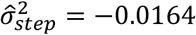 (upper panel) and 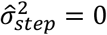 (lower panel), with maxima marked by circles with stars.

**Figure 9.**
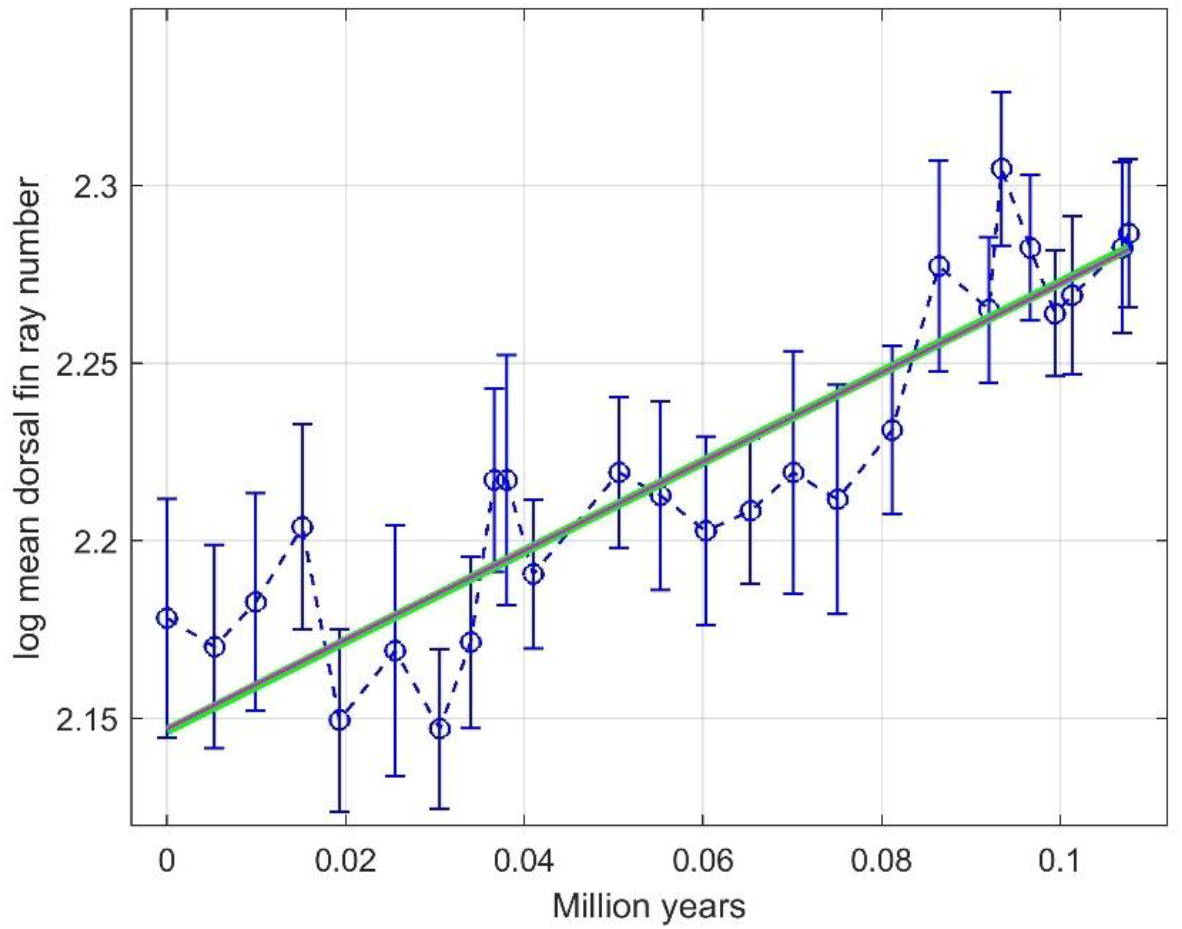
Results for the Stickleback fish case, with mean trait observations (dashed blue line with error bars), WLS predictions (green line), and GRW predictions with 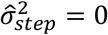 (red line, very much on top of the green line).

### 4.4 Stickleback fish case

In this final case I use data that show how the dorsal fin ray number evolved in the stickleback fish species *Gasterosteus doryssus* (Bell et al., 1985), see Table D4 in Appendix D for data. The zero point on the time scale is approximately −10 million years (personal information). In the data analysis the mean trait values are log-transformed, with the measurement errors transformed accordingly.

This is another example where the realistic step variance is found to be 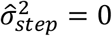, such that GLS is simplified into WLS. See results in Fig. 9 and Table 2.

As shown in Ergon (2025b), it is also in this case possible to find a tracking model. The essential elements and results for such a model are given in Appendix A.

## 5 Summary and discussion

The main aim of this paper is to show that the general random walk (GRW) model of Hunt (2006) with realistic measurement errors may be both under- and overestimating the degree of directional evolution, and that generalized least squares (GLS) estimation is a better alternative. A second aim is to show that realistically large measurement errors quite often result in a negative estimated step variance such that the step variance must be set to zero, with the consequence that the GRW model collapses into a deterministic directional walk model. A final aim is to show that adaptive peak tracking in some cases can be a good modeling alternative.

In the simulations, Hunt (2006) used regular sampling as explained in Section 2, and what appears to be the unrealistically low phenotypic variance *V*_*p*_ = 1, and one of the findings was that with 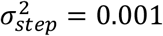 “some simulated sequences” yielded 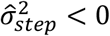 (Fig. 4 in Hunt, 2006). This is not surprising considering the term in Eq. 1 (with Hunt’s numbers) 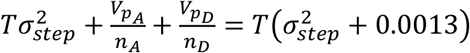, which obviously makes it rather difficult to estimate 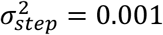 from data. Repeating the simulations in Hunt (2006) I found that “some” is around 10%, which with my simulations with 10 samples instead of 20 increases to 18%. With the more realistic phenotypic variance *V*_*p*_ = 400 and regular sampling I used 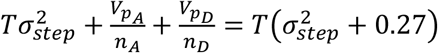, and for 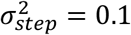 I found 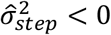 in 40% of the realizations (Table 1). The probability of obtaining 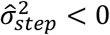 will thus increase when the number of samples decreases and the phenotypic variance increases. In the special case with step variance much larger than the step mean value, it is also difficult to estimate the step mean size, and it can in fact get the wrong sign (Appendix C).

In addition to simulations, different models were compared in four real data cases. In all these cases the estimated step variances were negative if they were allowed to be so, such that 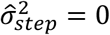 must be used in the GRW models, while GLS was simplified into WLS. In the Bryozoan and Stickleback fish cases I found that the GRW and WLS predictions were quite similar (Figures 4 and 9, and Table 2). In the Ostracod case 1 GRW underestimated the prediction slope by 21% (Fig. 5 and Table 2), while GRW in the Ostracod case 2 underestimated the prediction slope by 41% (Fig. 7 and Table 2).

The GRW model is not primarily intended for predictions, but once the mean step value is estimated a prediction model can be found by a generalized least squares approach. This makes it possible to compare GRW and GLS/WLS results using weighted mean squared errors (*WMSE*), and for all four of the real data cases with 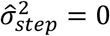 I found *WMSE*_*WLS*_ < *WMSE*_*GRW*_, although the difference is small in the Bryozoan case and very small in the Stickleback fish case (Table 2). This is not surprising, of course, since with zero step variance the GRW model collapses into a deterministic walk model, and there is no reason to believe that it then should perform better than a WLS model, which after all gives the best linear unbiased estimate (BLUE) of the evolutionary slope.

In three of the real data cases, I include an adaptive peak tracking model based on a dominating and well-known evolutionary driver (Ergon (2025a), and the tracking models gave better prediction results than GRW (Table 2). In the two Ostracod cases, however, the tracking results were poorer than the WLS results, which may possibly be explained by how the Mg/Ca temperature proxy was found. As shown in Ergon (2025b), a tracking model for Ostracod case 1 using the ∂^18^*O*(*t*) temperature proxy, gives clearly improved results. Ergon (2025b) also includes a tracking model for the stickleback fish case, but then by use of an association with rather than a direct measurement of an environmental proxy. Essential elements and results for such a model are given in Appendix A.

As shown in Table 1, *WMSE*_*GRW*_ ≈ *WMSE*_*GLS*_ in simulations with *V*_*p*_ = 1, while simulations with the more realistic phenotypic variance *V*_*p*_ = 400 show that *WMSE*_*GLS*_ < *WMSE*_*GRW*_. This is also the result in each of the four real data cases with 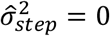, where GLS is replaced by WLS (Table 2), although the difference is marginal in two of the cases. The main conclusion is thus that GLS/WLS are the best methods for comparison of directional evolution with other evolutionary modes. As discussed in Hunt (2006), such comparisons are preferably performed using the Akaike Information Criterion (AIC), which raises questions of AIC for GLS and WLS. AIC for WLS is discussed in Ergon (2025b), based on Banks and Joyner (2017), but methods for finding AIC for GLS appear to be missing in the literature.

What this conclusion might mean for model categorization of fossil time series, as done in, e.g., Hunt et al. (2015) and Voje et al. (2018), is a question beyond the scope of this paper. It is, however, natural to ask how many of the time series in these studies that are examples of adaptive peak tracking as in the four real data cases in this paper. In such examples the different categories of random walk models may be of limited interest, since the evolutionary responses primarily are determined by the time series of the environmental drivers. The positive directional evolution in the Stickleback fish case, for example, would most likely have been negative if the fossils had been 0.2 million years younger (see Fig. B1 in Ergon, 2025b).

An interesting statistical issue for future research is the strong dependence on accurate estimates of sampling times. These are certainly measured with substantial errors, and the incremental change from one sample to another is a function of the time *T* between them. Hence, both the GRW and GLS/WLS solutions are influenced by errors in estimation of *T*, which possibly could cause large errors in the estimation results. This problem is discussed in Ergon (2025a) in relation to the Bryozoan case, but a more theoretical approach supported by a simulation study might be clarifying.

## Supporting information

MATAB Code

## Acknowledgments

I thank one of the reviewers for corrections regarding the covariance structure used in the simulations and for an interesting literature reference. I also thank University of South-Eastern Norway for support and funding.

## Conflicts of Interest

There are no conflicts of interest.

## Data Availability Statement

MATLAB code is archived on biorxiv https://doi.org/10.1101/2025.11.11.687792 Oxygen isotope data for Fig. 4 are archived as Raw data for simulations on bioRxiv, https://doi.org/10.1101/2024.10.30.621046.

## Appendix A. Tracking model for stickleback fish case

As shown in Ergon (2025b) it is possible to find a tracking model also in the stickleback fish case, with use of ∂^18^*O*(*t*) data as found from deep sea drilling samples (Westerhold et al., 2020). These data include obliquity cycles with a period of around 41,000 years, as found by power spectral density analysis (Fig. A1), and this information can be utilized by use of centered moving average smoothing as a feature extraction method. The result is shown in Fig. A2, from which follows *WMSE*_*WLS*_ = 0.000401 and *WMSE*_*tracking*_ = 0.000211.

The directional trend in Fig. A2 appears to be caused by an overall temperature trend, while the cycles apparently are caused by obliquity cycles. The model makes use of an association between cycles in the ∂^18^*O*(*t*) data and in the stickleback mean trait value, but the detailed mechanisms remain to be explained. It is for example well known that obliquity affects the intensity of seasons, and one mechanism may therefore be that this affects the balance between summer survival (declining with number of spines) and winter survival (increasing with number of spines), see Hendry et al. (2013).

**Figure A1.**
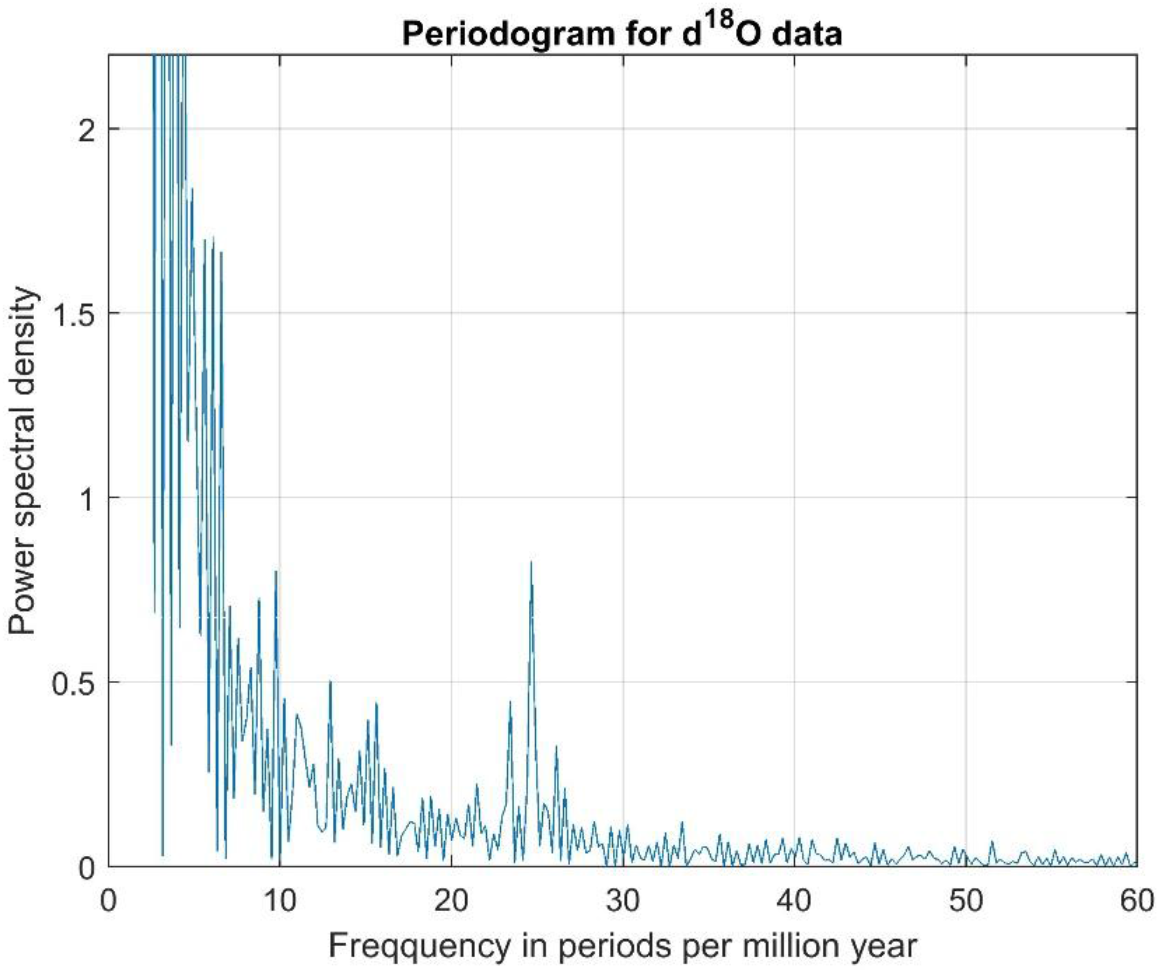
Periodogram for ∂^18^*O*(*t*), with obliquity cycles seen as a peak at 24.7 periods per million years.

**Figure A2.**
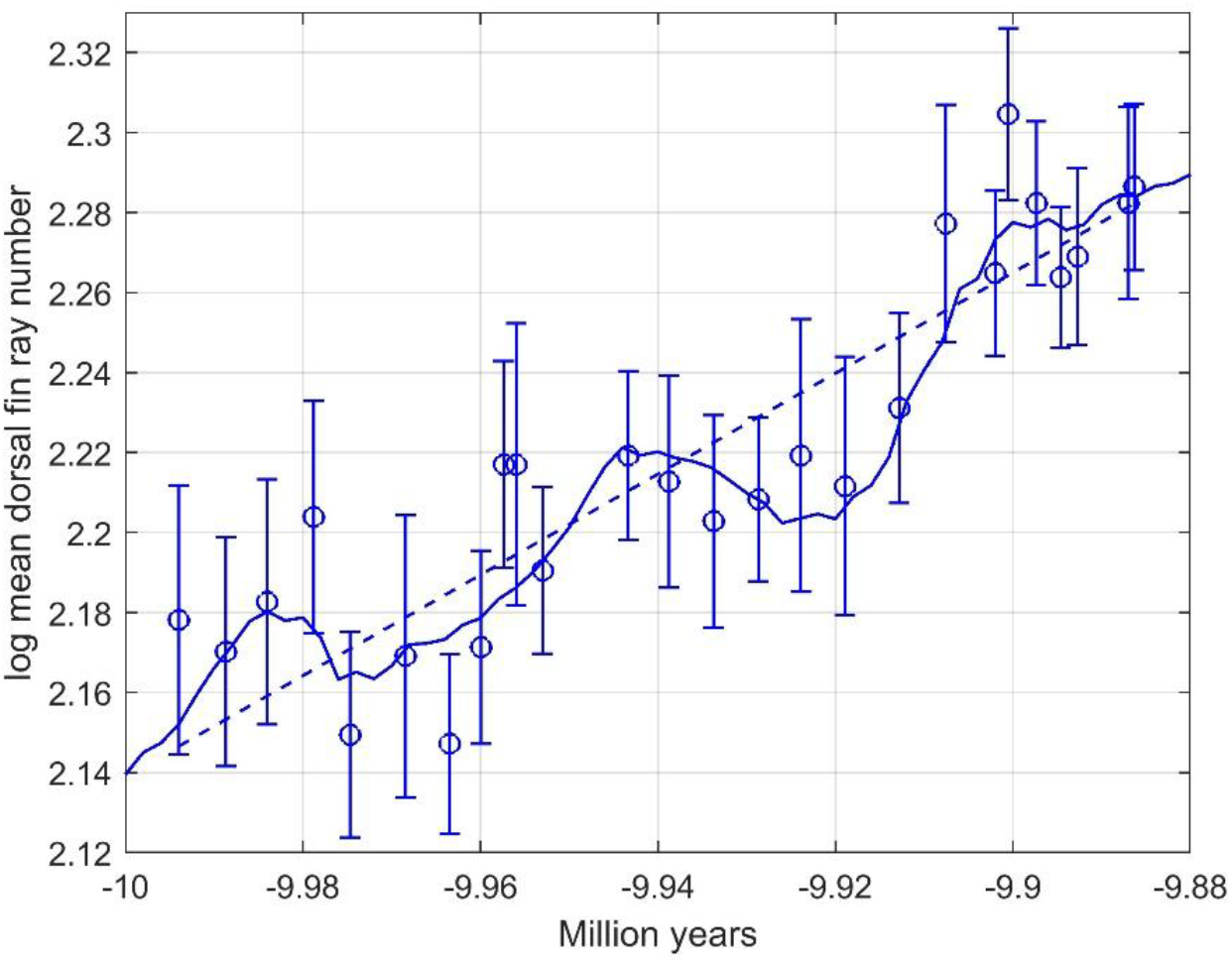
Predictions of stickleback fish mean trait by use of true data with obliquity cycles (solid line). Dashed line shows WLS predictions.

## Appendix B. Detailed simulation results

**Table B1.** Simulation 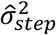 results with *V*_*p*_ = 1, based on 1,000 realizations with regular sampling and a fixed number of samples, *n* = 30. Numbers in parentheses are the number of realizations with 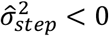.

| $\hat{\mu}_{step}$ | $\sigma_{step}^2 = 0.001$ | $\sigma_{step}^2 = 0.01$ | $\sigma_{step}^2 = 0.1$ | $\sigma_{step}^2 = 1$ | $\sigma_{step}^2 = 10$ |
| --- | --- | --- | --- | --- | --- |
| 0 | $0.00091 \pm 0.00082$<br>(120) | $0.0088 \pm 0.0047$<br>(0) | $0.090 \pm 0.045$<br>(0) | $0.88 \pm 0.45$<br>(0) | $8.7 \pm 4.4$<br>(0) |
| 0.01 | $0.00095 \pm 0.00080$<br>(91) | $0.0090 \pm 0.0048$<br>(0) | $0.088 \pm 0.044$<br>(0) | $0.87 \pm 0.42$<br>(0) | $9.0 \pm 4.3$<br>(0) |
| 0.1 | $0.00086 \pm 0.00075$<br>(102) | $0.0087 \pm 0.0047$<br>(1) | $0.090 \pm 0.045$<br>(0) | $0.91 \pm 0.45$<br>(0) | $9.1 \pm 4.5$<br>(0) |

**Table B2.** Simulation results with *V*_*p*_ = 400, based on 1,000 realizations with irregular sampling and variable numbers of samples, *n* = 3 to *n* = 60. Numbers in parentheses are the number of realizations with 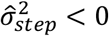.

| $\mu_{step}$ | $\sigma_{step}^2 = 0.001$ | $\sigma_{step}^2 = 0.01$ | $\sigma_{step}^2 = 0.1$ | $\sigma_{step}^2 = 1$ | $\sigma_{step}^2 = 10$ |
| --- | --- | --- | --- | --- | --- |
| 0 | $0.062 \pm 0.153$<br>(658) | $0.06 \pm 0.13$<br>(657) | $0.11 \pm 0.18$<br>(487) | $0.85 \pm 0.66$<br>(41) | $8.8 \pm 4.6$<br>(0) |
| 0.01 | $0.052 \pm 0.114$<br>(667) | $0.06 \pm 0.15$<br>(625) | $0.12 \pm 0.19$<br>(453) | $0.87 \pm 0.69$<br>(56) | $9.1 \pm 4.9$<br>(0) |
| 0.1 | $0.061 \pm 0.147$<br>(651) | $0.06 \pm 0.15$<br>(654) | $0.11 \pm 0.18$<br>(468) | $0.88 \pm 0.65$<br>(50) | $8.9 \pm 4.8$<br>(1) |

## Appendix C. Results for various values of 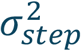

Fig. C1 shows histograms as in Fig. 2, but with 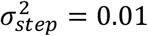, 0.1 and 1. Note that the variance of 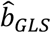 increases with 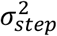, while the variance of the GRW estimation error 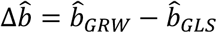 decreases.

**Figure C1.**
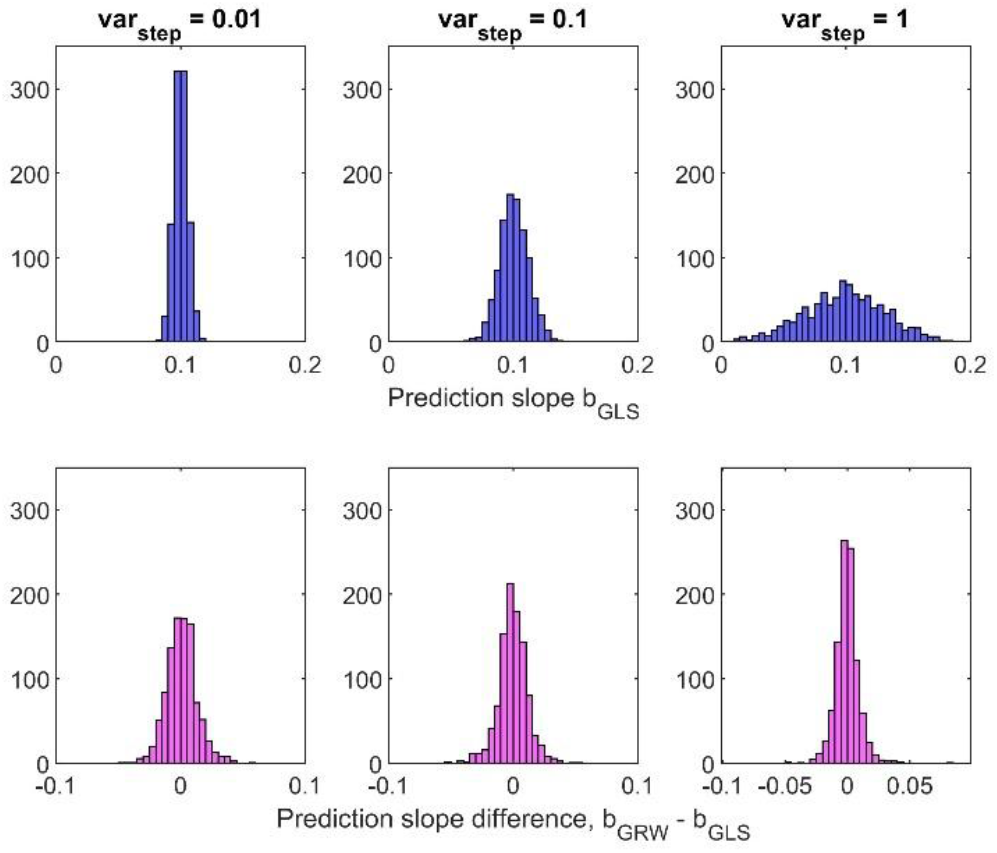
Typical histograms as in Fig. 2, for three values of 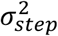.

As illustrations of the difficulties involved, Fig. C2 and Table C show responses and numerical results for two realizations of an evolutionary system with *µ*_*step*_ = 0.1 and 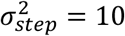, when the phenotypic variance is *V*_*p*_ = 400. The sampling was irregular. Note that the estimated slopes have different signs, and that the deviations from the GLS prediction lines are large compared to the standard errors. Also in these cases, however, *WMSE*_*GLS*_ is clearly lower than *WMSE*_*GRW*_.

**Figure C2.**
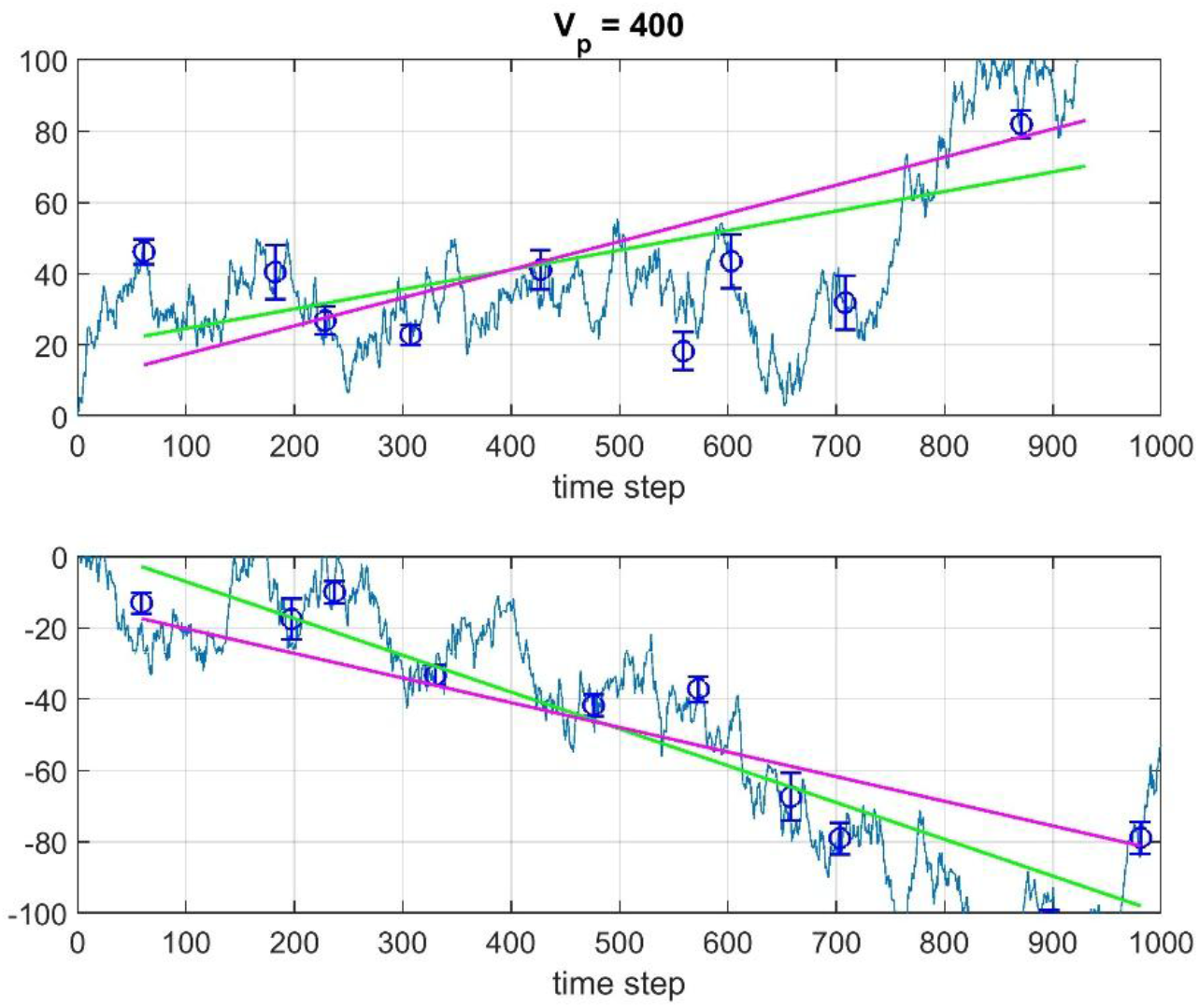
Responses for two realizations with *µ*_*step*_ = 0.1, 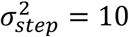, *V*_*p*_ = 400 and irregular sampling, with GLS (green lines) and GRW (red lines) predictions.

**Table C.** Estimated parameter values and WMSE results for responses in Fig. C2, i.e., for *µ*_*step*_ = 0.1, 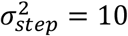, *V*_*p*_ = 400 and irregular sampling.

|  | Upper panel | Lower panel |
| --- | --- | --- |
| $\hat{\sigma}_{step}^2$ | 5.3 | 2.4 |
| $\hat{b}_{GRW} = \hat{\mu}_{step}$ | 0.08 | -0.07 |
| $\hat{b}_{WLS}$ | 0.05 | -0.10 |
| $WMSE_{GRW}$ | 309 | 194 |
| $WMSE_{GLS}$ | 271 | 124 |

## Appendix D Data for real data cases

**Table D1.**
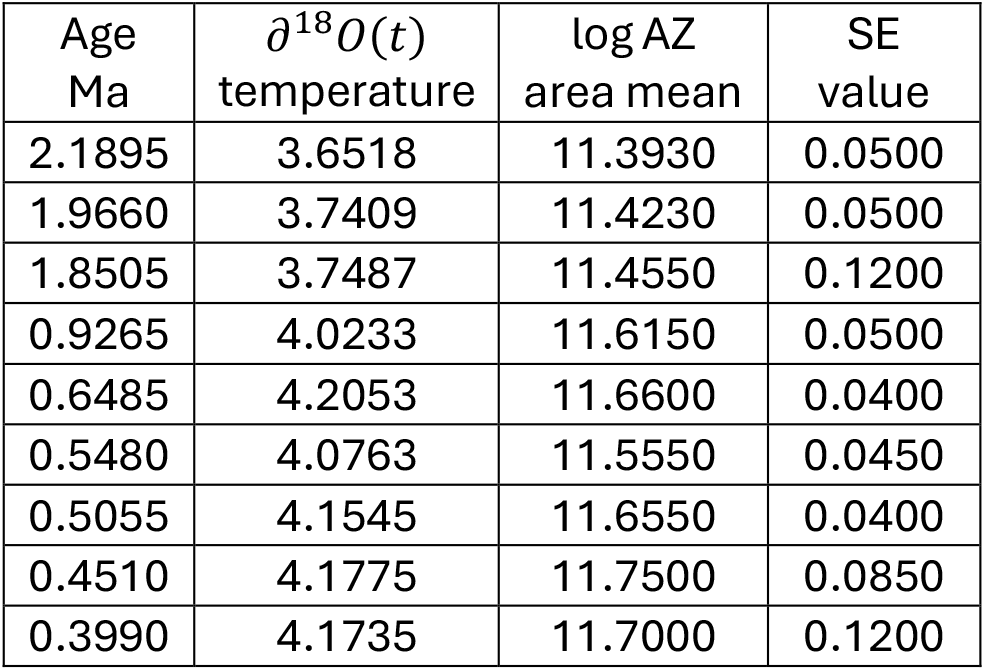
Data for Bryozoan case.

| Age<br>Ma | $\delta^{18}O(t)$<br>temperature | log AZ<br>area mean | SE<br>value |
| --- | --- | --- | --- |
| 2.1895 | 3.6518 | 11.3930 | 0.0500 |
| 1.9660 | 3.7409 | 11.4230 | 0.0500 |
| 1.8505 | 3.7487 | 11.4550 | 0.1200 |
| 0.9265 | 4.0233 | 11.6150 | 0.0500 |
| 0.6485 | 4.2053 | 11.6600 | 0.0400 |
| 0.5480 | 4.0763 | 11.5550 | 0.0450 |
| 0.5055 | 4.1545 | 11.6550 | 0.0400 |
| 0.4510 | 4.1775 | 11.7500 | 0.0850 |
| 0.3990 | 4.1735 | 11.7000 | 0.1200 |

**Table D2.**
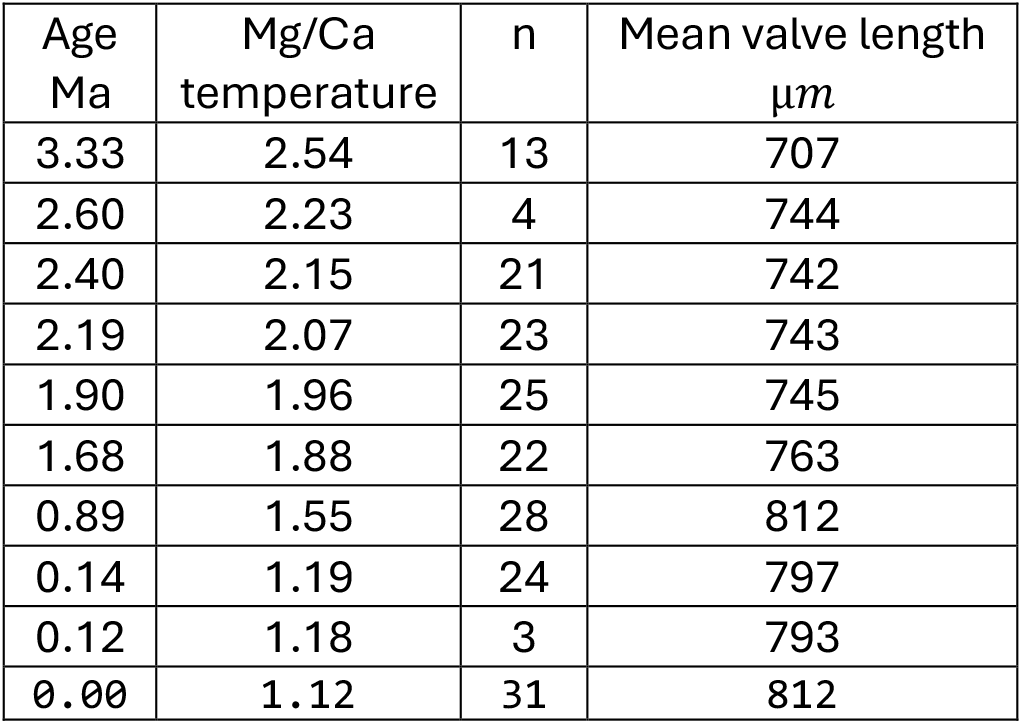
Data for Ostracode case 1.

**Table D3.** Data for Ostracode case 2.

| Age<br>Ma | Mg/Ca<br>temperature | n | Mean valve length<br>$\mu m$ |
| --- | --- | --- | --- |
| 3.96 | 2.79 | 13 | 616 |
| 3.33 | 2.54 | 1 | 681 |
| 3.22 | 2.49 | 7 | 734 |
| 2.58 | 2.21 | 6 | 704 |
| 2.19 | 2.07 | 1 | 708 |
| 1.68 | 1.88 | 2 | 723 |
| 0.21 | 1.22 | 5 | 776 |
| 0.11 | 1.17 | 29 | 777 |
| 0.09 | 1.17 | 43 | 767 |
| 0.08 | 1.16 | 35 | 770 |

**Table D4.**
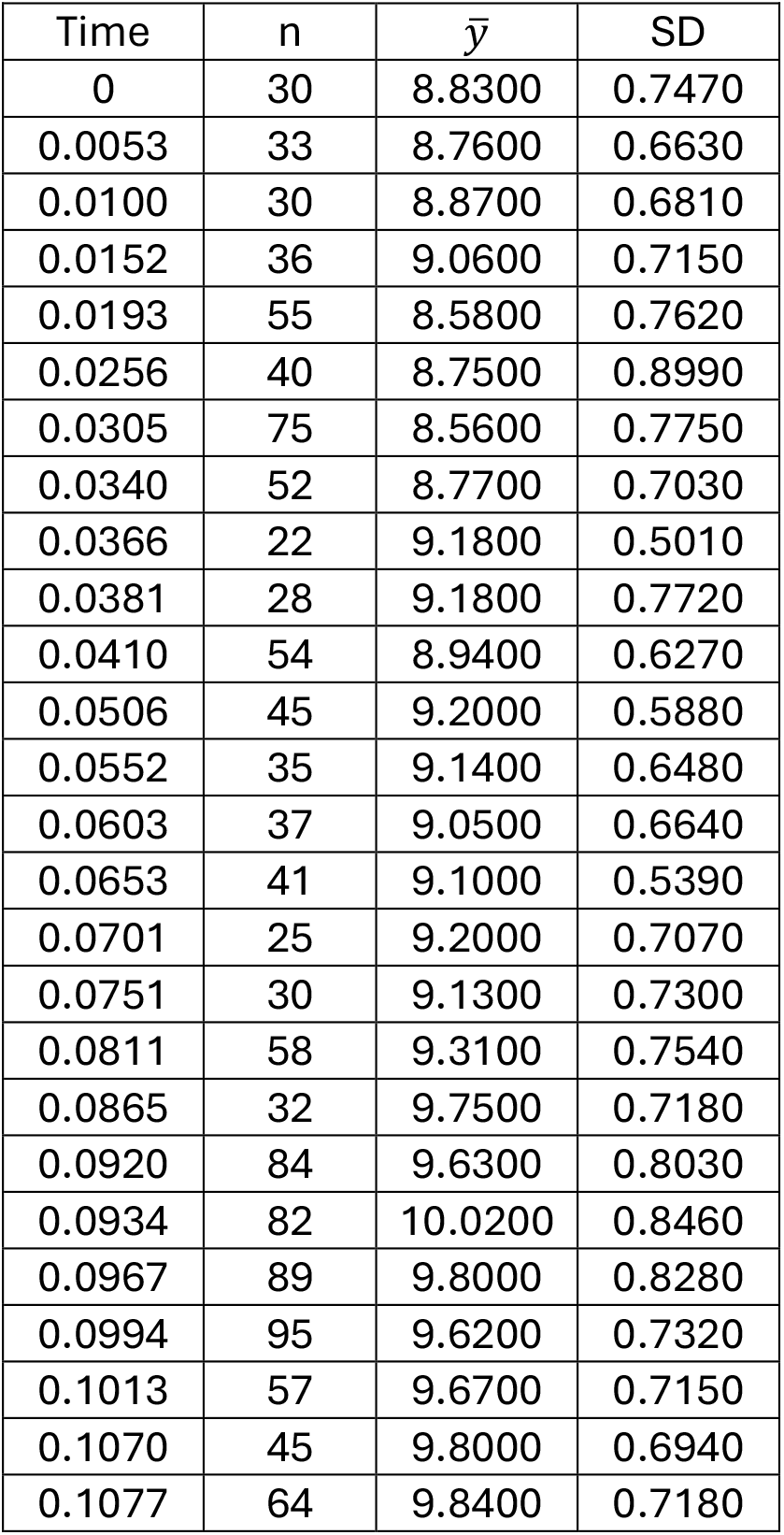
Data for Stickleback fish case.

## Notes

### Competing Interest Statement

The authors have declared no competing interest.

### Summary of Updates

The covariance function used in simulations is corrected. AIC comparisons are removed.

