## Supplementary material for "A critical look at directional random walk modeling of sparse fossil data": MATAB Code

MATLAB code for

Rolf Ergon

University of South-Eastern Norway

April 13, 2026

### Figures 1 and 3, and Table 1

```
clear

mu_step=0.1;
var_step=0.1;
Vp=400;
count=0;
y0=0;
M=100
for m=1:M
%% Generate data
y=y0*zeros(1,1000);
for t=2:1000
   tplot(t)=t;
   y(t)=y(t-1)+mu_step+sqrt(var_step)*randn;
end
tlabel=zeros(1,10);
ylabell=zeros(1,10);
err=zeros(1,10);

% j=1;                % Use for regular sampling
% for t=1:1000
%     if t/100==j
%        tlabel(j)=t-50;
```

```
%       j=j+1;
%    end
% end
% tlabel=round(tlabel);
% n=30*ones(1,10);

j=1;              % Use for irregular sampling
for j=1:10
   for t=1:1000
      if t/100==j
         tlabel(j)=t-99*rand;
         j=j+1;
      end
   end
end
tlabel=round(tlabel);
for j=1:10
   n(1,j)=57*rand;
end
n=round(n)+3*ones(1,10);

j=1;
for t=1:1000
   if t==tlabel(j)
      individuals=sqrt(Vp)*randn(n(j),1);
      Mean(1,j)=mean(individuals);
      ytrue(j)=y(t);
      ylabell(j)=y(t)+Mean(1,j);
      err(j)=sqrt(Vp/n(j));
      if j==10 break
      end
      j=j+1;
   end
end

%% Model 1: WLS on y data

tv=tlabel;
```

```
Cov0=[tv(1) tv(1) tv(1) tv(1) tv(1) tv(1) tv(1) tv(1) tv(1) tv(1)
    tv(1) tv(2) tv(2) tv(2) tv(2) tv(2) tv(2) tv(2) tv(2) tv(2)
    tv(1) tv(2) tv(3) tv(3) tv(3) tv(3) tv(3) tv(3) tv(3) tv(3)
    tv(1) tv(2) tv(3) tv(4) tv(4) tv(4) tv(4) tv(4) tv(4) tv(4)
    tv(1) tv(2) tv(3) tv(4) tv(5) tv(5) tv(5) tv(5) tv(5) tv(5)
    tv(1) tv(2) tv(3) tv(4) tv(4) tv(5) tv(6) tv(6) tv(6) tv(6)
    tv(1) tv(2) tv(3) tv(4) tv(5) tv(6) tv(7) tv(7) tv(7) tv(7)
    tv(1) tv(2) tv(3) tv(4) tv(5) tv(6) tv(7) tv(8) tv(8) tv(8)
    tv(1) tv(2) tv(3) tv(4) tv(5) tv(6) tv(7) tv(8) tv(9) tv(9)
    tv(1) tv(2) tv(3) tv(4) tv(5) tv(6) tv(7) tv(8) tv(9) tv(10)];

Cov1=Cov0*var_step;

v=err.^2;
V=diag(v);

X=[ones(10,1) tlabel'];
bls=inv(X'*inv(Cov1+V)*X)*X'*inv(Cov1+V)*ylabell';
a1=bls(1);
b1(m)=bls(2);
for i=1:10
   yhat1(i)=a1+b1(m)*tlabel(i);
end
WMSE_GLS(m)=(ytrue-yhat1)*inv(Cov1+V)*(ytrue-yhat1)'/trace(inv(Cov1+V));

%% Model 2: GRW
n_samples=ones(1,10);
var_samples=err.^2;

c=1;
Tsample=c*tlabel;

for i=1:9
   dT(i)=-Tsample(i)+Tsample(i+1);
   dX(i)=ylabell(i+1)-ylabell(i);
   nA(i)=n_samples(i);
   nD(i)=n_samples(i+1);
```

```
    varA(i)=var_samples(i);
    varD(i)=var_samples(i+1);
end

% Contraints
mustep_min=-100; mustep_max=100;
varstep_min=-100; varstep_max=100;
par_lb=[mustep_min varstep_min];
par_ub=[mustep_max varstep_max];

% fmincon
par_guess=[0 0];
Aineq=[]; Bineq=[]; Aeq=[]; Beq=[];
fun_objective_handle=...
    @(par)fun_objective(par,dT,dX,varA,nA,varD,nD);
[par_opt,fval,exitflag,output,lambda,grad,hessian] =...
fmincon(fun_objective_handle,par_guess,Aineq,Bineq,Aeq,Beq,par_lb,par_ub);

mustep(m)=par_opt(1);
varstep(m)=par_opt(2);

if varstep(m)<0
    count=count+1;
end

varstepestimated(m)=varstep(m);

if varstep(m)<0
    varstep(m)=0;
end

Cov2=Cov0*varstep(m);

X=ones(10,1);
b2=c*mustep(m);
a2=inv(X'*inv(Cov2+V)*X)*X'*inv(Cov2+V)*(ylabell'-b2*tlabel');
for j=1:10
    yhat2(j)=a2+b2*tlabel(j);
```

```
end

WMSE_GRW(m)=sum((ytrue-yhat2)*inv(Cov2+V)*(ytrue-yhat2)')/trace(inv(Cov2+V));

ratio(m)=WMSE_GLS(m)/WMSE_GRW(m);

end

Mstep=[mean(mustep) std(mustep)]
Varstep=[mean(varstep) std(varstep)]
B1=[mean(b1) std(b1)]
WMSEGRW=[mean(WMSE_GRW) std(WMSE_GRW)]
WMSEGLS=[mean(WMSE_GLS) std(WMSE_GLS)]
count

figure(1)
subplot(1,2,2)
plot(tplot,y), hold on
errorbar(tlabel,ylabell,err,'bo','LineWidth',1.0), hold on
plot(tlabel,yhat1,'g','LineWidth',1.0)
plot(tlabel,yhat2,'m','LineWidth',1.0)
hold off, grid
xlabel('time step')
title('V_p = 1')
axis([0 1000 0 100])
% axis([0 1000 -100 0])

%% Parameter search
function f = fun_objective(par,dT,dX,varA,nA,varD,nD)
mustep=par(1);
varstep=par(2);

for i=1:9
  varterm(i)=dT(i)*varstep+varA(i)/nA(i)+varD(i)/nD(i);
  Li(i)=-log(2*pi)/2-log(varterm(i))/2-(dX(i)-dT(i)*mustep)^2/(2*varterm(i));
end

L=sum(Li);
```

```
f=-L;


end
```

### Figure 2

```
clear


mu_step=0.1;
var_step=0.1;
Vp=400;
count=0;
y0=0;
M=1000


for m=1:M
%% Generate data
y=y0*zeros(1,1000);
for t=2:1000
   tplot(t)=t;
   y(t)=y(t-1)+mu_step+sqrt(var_step)*randn;
end
tlabel=zeros(1,10);
ylabell=zeros(1,10);
err=zeros(1,10);


j=1;
for j=1:10
   for t=1:1000
      if t/100==j
         tlabel(j)=t-99*rand;
         j=j+1;
      end
   end
end
tlabel=round(tlabel);
for j=1:10
```

```
   n(1,j)=57*rand;
end
n=round(n)+3*ones(1,10);


j=1;
for t=1:1000
   if t==tlabel(j)
      individuals=sqrt(Vp)*randn(n(j),1);
      Mean(1,j)=mean(individuals);
      ytrue(j)=y(t);
      ylabell(j)=y(t)+Mean(1,j);
      err(j)=sqrt(Vp/n(j));
      if j==10 break
      end
      j=j+1;
   end
end

%% Model 1: WLS on y data

tv=tlabel;

Cov0=[tv(1) tv(1) tv(1) tv(1) tv(1) tv(1) tv(1) tv(1) tv(1) tv(1)
    tv(1) tv(2) tv(2) tv(2) tv(2) tv(2) tv(2) tv(2) tv(2)
    tv(1) tv(2) tv(3) tv(3) tv(3) tv(3) tv(3) tv(3) tv(3) tv(3)
    tv(1) tv(2) tv(3) tv(4) tv(4) tv(4) tv(4) tv(4) tv(4) tv(4)
    tv(1) tv(2) tv(3) tv(4) tv(5) tv(5) tv(5) tv(5) tv(5) tv(5)
    tv(1) tv(2) tv(3) tv(4) tv(4) tv(5) tv(6) tv(6) tv(6) tv(6)
    tv(1) tv(2) tv(3) tv(4) tv(5) tv(6) tv(7) tv(7) tv(7) tv(7)
    tv(1) tv(2) tv(3) tv(4) tv(5) tv(6) tv(7) tv(8) tv(8) tv(8)
    tv(1) tv(2) tv(3) tv(4) tv(5) tv(6) tv(7) tv(8) tv(9) tv(9)
    tv(1) tv(2) tv(3) tv(4) tv(5) tv(6) tv(7) tv(8) tv(9) tv(10)];

Cov1=Cov0*var_step;

v=err.^2;
V=diag(v);
```

```
X=[ones(10,1) tlabel'];
bls=inv(X'*inv(Cov1+V)*X)*X'*inv(Cov1+V)*ylabell';
a1=bls(1);
b1(m)=bls(2);
for i=1:10
   yhat1(i)=a1+b1(m)*tlabel(i);
end
WMSE_GLS(m)=(ytrue-yhat1)*inv(Cov1+V)*(ytrue-yhat1)'/trace(inv(Cov1+V));

%% Model 2: GRW


n_samples=ones(1,10);
var_samples=err.^2;


c=1;
Tsample=c*tlabel;


for i=1:9
   dT(i)=-Tsample(i)+Tsample(i+1);
   dX(i)=ylabell(i+1)-ylabell(i);
   nA(i)=n_samples(i);
   nD(i)=n_samples(i+1);
   varA(i)=var_samples(i);
   varD(i)=var_samples(i+1);
end

% Contraints
mustep_min=-100; mustep_max=100;
varstep_min=-100; varstep_max=100;
par_lb=[mustep_min varstep_min];
par_ub=[mustep_max varstep_max];

% fmincon
par_guess=[0 0];
Aineq=[]; Bineq=[]; Aeq=[]; Beq=[];
fun_objective_handle=...
   @(par)fun_objective(par,dT,dX,varA,nA,varD,nD);
```

```
[par_opt,fval,exitflag,output,lambda,grad,hessian] =...
fmincon(fun_objective_handle,par_guess,Aineq,Bineq,Aeq,Beq,par_lb,par_ub);

mustep(m)=par_opt(1);
varstep(m)=par_opt(2);

if varstep(m)<0
   count=count+1;
end

varstepestimated(m)=varstep(m);

if varstep(m)<0
   varstep(m)=0;
end

Cov2=Cov0*varstep(m);

X=ones(10,1);
b2=c*mustep(m);
a2=inv(X'*inv(Cov2+V)*X)*X'*inv(Cov2+V)*(ylabell'-b2*tlabel');
for j=1:10
   yhat2(j)=a2+b2*tlabel(j);
end

WMSE_GRW(m)=sum((ytrue-yhat2)*inv(Cov2+V)*(ytrue-yhat2)')/trace(inv(Cov2+V));

ratio(m)=WMSE_GLS(m)/WMSE_GRW(m);

end

count

figure(2)
subplot(2,1,1)
histogram(b1,40,'BinWidth',0.005,'FaceColor','b')
ax = findobj(subplot(2,1,1),'Type','Axes');
for i = 1:length(ax)
```

```
    ylim(ax(i),[0 200]);
    xlim(ax(i),[0.05 0.15]);
end
xlabel('Prediction slope b_G_L_S')
subplot(2,1,2)
histogram(mustep-b1,40,'BinWidth',0.005,'FaceColor','m')
ax = findobj(subplot(2,1,2),'Type','Axes');
for i = 1:length(ax)
    ylim(ax(i),[0 200]);
    xlim(ax(i),[-0.05 0.05]);
end
xlabel('Prediction slope difference, b_G_R_W - b_G_L_S')

%% Parameter search
function f = fun_objective(par,dT,dX,varA,nA,varD,nD)
mustep=par(1);
varstep=par(2);

for i=1:9
    varterm(i)=dT(i)*varstep+varA(i)/nA(i)+varD(i)/nD(i);
    Li(i)=-log(2*pi)/2-log(varterm(i))/2-(dX(i)-dT(i)*mustep)^2/(2*varterm(i));
end

L=sum(Li);

f=-L;

end
```

### Figure 4

```
clear
window=100;

%% Load O18 data, order and plot
load O18.mat
x=5320-x;
x=flipud(x);
```

```
u=flipud(y);

x=0.001*x;              % Million year scale
ufilt=movmean(u,window);
x=x(737:2115);
u=u(737:2115);
ufilt=ufilt(737:2115);
tplot=x-5.3-0.02;         % -0.2 for zero point correction

% Sample data from Liow et al. (2024)
tlabel=[-2.1895 -1.9660 -1.8505 -0.9265 -0.6485 -0.5480 -0.5055 -0.4510 -0.3990 ];
ylabell=[11.393 11.423 11.455 11.615 11.66 11.555 11.655 11.75 11.7 ];
err=[0.05 0.05 0.12 0.05 0.04 0.045 0.04 0.085 0.12];

%% Tracking model
ulabel=[mean(ufilt(54))
    mean(ufilt(143))
    mean(ufilt(189))
    mean(ufilt(616))
    mean(ufilt(755))
    mean(ufilt(831))
    mean(ufilt(874))
    mean(ufilt(928))
    mean(ufilt(980))
    ]';
w=err.^-2;
W=diag(w);
W1=W;
X=[ones(9,1) ulabel'-3.6*ones(9,1)];
bls=inv(X'*W1*X)*X'*W1*ylabell';
ahat1=bls(1);
bhat1=bls(2);

for t=1:1379
   yhatplot(t)=ahat1+bhat1*(ufilt(t)-3.6);
end
yhat1=ahat1+bhat1*(ulabel-3.6*ones(1,9));
```

```
WMSE_Tracking=sum((ylabell-yhat1)*W1*(ylabell-yhat1)')/sum(diag(W1));

yplot=bls(1)+bls(2)*(ulabel'-3.6*ones(9,1));

%% WLS model
W2=W1;

X=[ones(9,1) tlabel'];
bls=inv(X'*W2*X)*X'*W2*ylabell';
ahat2=bls(1);
bhat2=bls(2);
for i=1:9
  yhat2(1,i)=ahat2+bhat2*tlabel(i);
end

WMSE_WLS=sum((ylabell-yhat2)*W2*(ylabell-yhat2)')/sum(diag(W2));

%% GRW model
W3=W2;

n_samples=ones(1,10);
var_samples=err.^2;

c=1;
Tsample=c*tlabel;

for i=1:8
  dT(i)=-Tsample(i)+Tsample(i+1);
  dX(i)=ylabell(i+1)-ylabell(i);
  nA(i)=n_samples(i);
  nD(i)=n_samples(i+1);
  varA(i)=var_samples(i);
  varD(i)=var_samples(i+1);
end

% Contraints
mustep_min=-1; mustep_max=1;
varstep_min=0; varstep_max=1;
```

```
par_lb=[mustep_min varstep_min];

par_ub=[mustep_max varstep_max];


par_guess=[0 0];

Aineq=[]; Bineq=[]; Aeq=[]; Beq=[];

fun_objective_handle=...

   @(par)fun_objective(par,dT,dX,varA,nA,varD,nD);

[par_opt,fval,exitflag,output,lambda,grad,hessian] =...

fmincon(fun_objective_handle,par_guess,Aineq,Bineq,Aeq,Beq,par_lb,par_ub);


par_opt;

varstep=par_opt(2)

mustep=par_opt(1)


X=ones(9,1);

bhat3=c*mustep;

ahat3=inv(X'*W3*X)*X'*W3*(ylabell'-bhat3*tlabel');

for i=1:9

   yhat3(i)=ahat3+bhat3*tlabel(i);

end


WMSE_GRW=sum((ylabell-yhat3)*W3*(ylabell-yhat3)')/sum(diag(W3));


bhat2

bhat3

WMSE_Tracking

WMSE_WLS

WMSE_GRW


figure(4)

errorbar(tlabel,ylabell,err,'bo','LineWidth',1.0), hold on

errorbar(tlabel,ylabell,err,'--b','LineWidth',1.0)

plot(tplot,yhatplot,'b','LineWidth',1.0)

plot(tlabel,yhat2,'g','LineWidth',1.0)

plot(tlabel,yhat3,'m','LineWidth',1.0)

hold off, grid

axis([-2.35 -0.3 11.3 11.85])
```

```
xlabel('Million years')
ylabel('log AZ area mean')

%% Parameter search
function f = fun_objective(par,dT,dX,varA,nA,varD,nD)
mustep=par(1);
varstep=par(2);

for i=1:8
   varterm(i)=dT(i)*varstep+varA(i)/nA(i)+varD(i)/nD(i);
   Li(i)=-log(2*pi)/2-log(varterm(i))/2-(dX(i)-dT(i)*mustep)^2/(2*varterm(i));
end

L=-sum(Li);

f=L;

end
```

### Figures 5 and 6

```
clear
s2=467;                 % micrometer^2

%% Sample data
Data=[
3.33 1,962 2.54 13 707
2.60 3,427 2.23 4 744
2.40 1,253 2.15 21 742
2.19 1,253 2.07 23 743
1.90 1,253 1.96 25 745
1.68 1,253 1.88 22 763
0.89 2,873 1.55 28 812
0.14 2,873 1.19 24 797
0.12 3,427 1.18 3 793
0.00 2,781 1.12 31 812
];
```

```
tlabel=-Data(:,1)';

y=Data(:,6)';

ylabell=log(Data(:,6))';

n=Data(:,5)';


for i=1:10

   err(i)=sqrt(s2/n(i))*ylabell(i)/y(i);

end


%% Tracking model

u=Data(:,4)';

w=err.^-2;

W1=diag(w);


X=[ones(10,1) u'-2.54*ones(10,1)];

bls=inv(X'*W1*X)*X'*W1*ylabell'

a=bls(1);

b=bls(2);


for i=1:10

   yhat1(i)=a+b*(u(i)-2.54);

end


WMSE_Tracking=sum((ylabell-yhat1)*W1*(ylabell-yhat1)')/sum(diag(W1));


yplot=bls(1)+bls(2)*(u'-2.54*ones(10,1));


%% WLS model

W2=W1;


X=[ones(10,1) tlabel'];

bls=inv(X'*W2*X)*X'*W2*ylabell';

a2=bls(1);

b2=bls(2);

for i=1:10

   yhat2(i)=a2+b2*tlabel(i);

end
```

```
WMSE_WLS=(ylabell-yhat2)*W2*(ylabell-yhat2)'/sum(diag(W2));

%% GRW model
W3=W2;
n_samples=ones(1,10);
var_samples=err.^2;

c=1;
Tsample=c*tlabel;

for i=1:9
   dT(i)=-Tsample(i)+Tsample(i+1);
   dX(i)=ylabell(i+1)-ylabell(i);
   nA(i)=n_samples(i);
   nD(i)=n_samples(i+1);
   varA(i)=var_samples(i);
   varD(i)=var_samples(i+1);
end

% Contraints
mustep_min=-1; mustep_max=1;
varstep_min=0; varstep_max=1;

varstep_min=-1; varstep_max=1;

par_lb=[mustep_min varstep_min];
par_ub=[mustep_max varstep_max];

par_guess=[0 0];
Aineq=[]; Bineq=[]; Aeq=[]; Beq=[];
fun_objective_handle=...
   @(par)fun_objective(par,dT,dX,varA,nA,varD,nD);
[par_opt,fval,exitflag,output,lambda,grad,hessian] =...
fmincon(fun_objective_handle,par_guess,Aineq,Bineq,Aeq,Beq,par_lb,par_ub);
output;

mustep=par_opt(1);
varstep=par_opt(2);
```

```
X=ones(10,1);
b3=c*mustep;
a3=inv(X'*W3*X)*X'*W3*(ylabell'-b3*tlabel');
for i=1:10
   yhat3(i)=a3+b3*tlabel(i);
end

WMSE_GRW=sum((ylabell-yhat3)*W3*(ylabell-yhat3)')/sum(diag(W3));

figure(5)
errorbar(tlabel,ylabell,err,'bo','LineWidth',1.0), hold on
errorbar(tlabel,ylabell,err,'--b','LineWidth',1.0)
plot(tlabel,yhat1,'b','LineWidth',1.0)
plot(tlabel,yhat2,'g','LineWidth',1.0)
plot(tlabel,yhat3,'m','LineWidth',1.0), hold off, grid
axis([-3.4 0.1 6.5 6.8])
xlabel('Million years')
ylabel('log mean body size')

mustep
b2
WMSE_Tracking
WMSE_WLS
WMSE_GRW

%% Grid
varstepg=1.25*varstep;
mustepg=0.5*mustep;
for q=1:100
   varstepg(q+1)=varstepg(q)+0.01e-3;
   for j=1:100
      mustepg(j+1)=mustepg(j)+0.01*mustep;
      for i=1:9
         vartermg(i)=dT(i)*varstepg(q)+varA(i)/nA(i)+varD(i)/nD(i);
         Lgi(i)=-log(2*pi)/2-log(vartermg(i))/2-(dX(i)-dT(i)*mustepg(j))^2/(2*vartermg(i));
      end
      Lg=sum(Lgi);
```

```
    zg(j,q)=-Lg;
  end
end

figure(6)
contour(varstepg(2:101),mustepg(2:101),zg,100), hold on
plot(varstep,mustep,'o')
plot(varstep,mustep,'*'),hold off, grid
xlabel('var_s_t_e_p')
ylabel('m_s_t_e_p')

%% Parameter search
function f = fun_objective(par,dT,dX,varA,nA,varD,nD)
mustep=par(1);
varstep=par(2);

for i=1:9
  varterm=dT(i)*varstep+varA(i)/nA(i)+varD(i)/nD(i);
  Li(i)=-log(2*pi)/2-log(varterm)/2-(dX(i)-dT(i)*mustep)^2/(2*varterm);
end

L=sum(Li);

f=-L

end
```

### Figures 7 and 8

```
clear
s2=467;          % micrometer^2

%% Sample data
Data=[
3.96 1,313 2.79 13 616
3.33 1,962 2.54 1 681
3.22 2,417 2.49 7 734
2.58 3,427 2.21 6 704
```

```
2.19 1,253 2.07 1 708
1.68 1,253 1.88 2 723
0.21 3,427 1.22 5 776
0.11 3,427 1.17 29 777
0.09 3,427 1.17 43 767
0.08 3,427 1.16 35 770
];

tlabel=-Data(:,1)';
y=Data(:,6)';
ylabell=log(Data(:,6))';
n=Data(:,5)';

for i=1:10
   err(i)=sqrt(s2/n(i))*ylabell(i)/y(i);
end

%% Tracking model
u=Data(:,4)';
w=err.^-2;
W1=diag(w);

X=[ones(10,1) u'-2.54*ones(10,1)];
bls=inv(X'*W1*X)*X'*W1*ylabell'
a=bls(1);
b=bls(2);

for i=1:10
   yhat1(i)=a+b*(u(i)-2.54);
end

WMSE_Tracking=sum((ylabell-yhat1)*W1*(ylabell-yhat1)')/sum(diag(W1));

%% WLS model
W2=W1;

X=[ones(10,1) tlabel'];
bls=inv(X'*W2*X)*X'*W2*ylabell';
```

```
a2=bls(1);
b2=bls(2);
for i=1:10
   yhat2(i)=a2+b2*tlabel(i);
end

WMSE_WLS=(ylabell-yhat2)*W2*(ylabell-yhat2)'/sum(diag(W2));

%% GRW model
n_samples=ones(1,10);
var_samples=err.^2;

c=1;
Tsample=c*tlabel;

for i=1:9
   dT(i)=-Tsample(i)+Tsample(i+1);
   dX(i)=ylabell(i+1)-ylabell(i);
   nA(i)=n_samples(i);
   nD(i)=n_samples(i+1);
   varA(i)=var_samples(i);
   varD(i)=var_samples(i+1);
end

% Contraints
mustep_min=-1; mustep_max=1;
varstep_min=0; varstep_max=1;
par_lb=[mustep_min varstep_min];
par_ub=[mustep_max varstep_max];

% fmincon
par_guess=[0 0];
Aineq=[]; Bineq=[]; Aeq=[]; Beq=[];
fun_objective_handle=...
   @(par)fun_objective(par,dT,dX,varA,nA,varD,nD);
[par_opt,fval,exitflag,output,lambda,grad,hessian] =...
fmincon(fun_objective_handle,par_guess,Aineq,Bineq,Aeq,Beq,par_lb,par_ub);
output;
```

```
mustep=par_opt(1);
varstep=par_opt(2);

for i=1:9
   varterm(i)=dT(i)*varstep+varA(i)/nA(i)+varD(i)/nD(i);
   Li(i)=-log(2*pi)/2-log(varterm(i))/2-(dX(i)-dT(i)*mustep)^2/(2*varterm(i));
end

L=sum(Li);

W3=W2;
X=ones(10,1);
b3=c*mustep;
a3=inv(X'*W3*X)*X'*W3*(ylabell'-b3*tlabel');
for i=1:10
   yhat3(i)=a3+b3*tlabel(i);
end

WMSE_GRW=sum((ylabell-yhat3)*W3*(ylabell-yhat3)')/sum(diag(W3));

figure(7)
errorbar(tlabel,ylabell,err,'bo','LineWidth',1.0), hold on
errorbar(tlabel,ylabell,err,'--b','LineWidth',1.0)
plot(tlabel,yhat1,'b','LineWidth',1.0)
plot(tlabel,yhat2,'g','LineWidth',1.0)
plot(tlabel,yhat3,'m','LineWidth',1.0)
hold off, grid
axis([-4.1 0 6.3 6.8])
xlabel('Million years')
ylabel('log mean body size')

mustep
b2
b3
WMSE_Tracking
WMSE_WLS
WMSE_GRW
```

```
%% Plot mu
varstepplot=varstep;
mustepplot=0.1e-2;
for j=1:100
  mustepplot(j+1)=mustepplot(j)+0.05e-2;
    for i=1:9
      vartermplot(i)=dT(i)*varstepplot+varA(i)/nA(i)+varD(i)/nD(i);
      Lgi(i)=-log(2*pi)/2-log(vartermplot(i))/2-(dX(i)-dT(i)*mustepplot(j))^2/(2*vartermplot(i));
    end
    Lg=sum(Lgi);
    Lplot(j)=Lg;
end

figure(8)
subplot(2,1,2)                    % Use for varstep > 0
plot(mustepplot(2:101),Lplot), hold on, grid
plot(mustep,L,'bo')
plot(mustep,L,'b*'), hold off
axis([0 5e-2 10 10.1])
xlabel('mu_s_t_e_p')
ylabel('log likelihood')
%
% subplot(2,1,1)                    % Use for varstep as free varaible
% plot(mustepplot(2:101),Lplot),hold on, grid
% plot(mustep,L,'bo')
% plot(mustep,L,'b*'), hold off
% axis([0 5e-2 11.2 11.8])
% ylabel('log likelihood')

%% Parameter search
function f = fun_objective(par,dT,dX,varA,nA,varD,nD)
mustep=par(1);
varstep=par(2);

for i=1:9
  varterm(i)=dT(i)*varstep+varA(i)/nA(i)+varD(i)/nD(i);
  Li(i)=-log(2*pi)/2-log(varterm(i))/2-(dX(i)-dT(i)*mustep)^2/(2*varterm(i));
```

```
end

L=sum(Li);

f=-L;

end
```

### Figure 9

```
clear

%% Sample data

tlabel1=[0 5317 9950 15182 19318 25551 30541 34030 36631 38052 41008 50629 55246];
tlabel2=[60289 65312 70119 75075 81150 86454 92006 93445 96659 99377 101306 106969 107667];
tlabel=[tlabel1 tlabel2]/1000000;
n=[30 33 30 36 55 40 75 52 22 28 54 45 35 37 41 25 30 58 32 84 82 89 95 57 45 64];
y1=0.01*[883 876 887 906 858 875 856 877 918 918 894 920 914 ];
y2=0.01*[905 910 920 913 931 975 963 1002 980 962 967 980 984];
y=[y1 y2];
sd1=0.001*[747 663 681 715 762 899 775 703 501 772 627 588 648  ];
sd2=0.001*[664 539 707 730 754 718 803 846 828 732 715 694 718];
sd=[sd1 sd2];
var=sd.^2;
err0=sqrt(var./n);

ylabell=log(y);

for i=1:26
   err(i)=err0(i)*(ylabell(i)/y(i))^1;
end

%% WLS model
w=err.^-2;
W2=diag(w);
```

```
X=[ones(26,1) tlabel'];
bls=inv(X'*W2*X)*X'*W2*ylabell';
a2=bls(1);
b2=bls(2);
for i=1:26
   yhat2(i)=a2+b2*tlabel(i);
end

WMSE_WLS=(ylabell-yhat2)*W2*(ylabell-yhat2)'/sum(diag(W2));

%% GRW model
W3=diag(w);
n_samples=ones(1,26);
var_samples=err.^2;

c=1;
Tsample=c*tlabel;

for i=1:25
   dT(i)=-Tsample(i)+Tsample(i+1);
   dX(i)=ylabell(i+1)-ylabell(i);
   nA(i)=n_samples(i);
   nD(i)=n_samples(i+1);
   varA(i)=var_samples(i);
   varD(i)=var_samples(i+1);
end

% Contraints
mustep_min=-100; mustep_max=100;
varstep_min=0; varstep_max=100;

par_lb=[mustep_min varstep_min];
par_ub=[mustep_max varstep_max];

par_guess=[0 0];
Aineq=[]; Bineq=[]; Aeq=[]; Beq=[];
fun_objective_handle=...
   @(par)fun_objective(par,dT,dX,varA,nA,varD,nD);
```

```
[par_opt,fval,exitflag,output,lambda,grad,hessian] =...
fmincon(fun_objective_handle,par_guess,Aineq,Bineq,Aeq,Beq,par_lb,par_ub);
output;

mustep=par_opt(1);
varstep=par_opt(2);

X=ones(26,1);
b3=c*mustep;
a3=inv(X'*W3*X)*X'*W3*(ylabell'-b3*tlabel');
for i=1:26
   yhat3(i)=a3+b3*tlabel(i);
end

WMSE_GRW=sum((ylabell-yhat3)*W3*(ylabell-yhat3)')/sum(diag(W3));


figure(9)
errorbar(tlabel,ylabell,err,'bo','LineWidth',1.0), hold on
errorbar(tlabel,ylabell,err,'--b','LineWidth',1.0)
plot(tlabel,yhat2,'g','LineWidth',3.0)
plot(tlabel,yhat3,'m','LineWidth',1.0)

hold off, grid
axis([-0.004 0.112 2.12 2.35])

xlabel('Million years')
ylabel('log mean dorsal fin ray number')

mustep
b2
WMSE_WLS
WMSE_GRW

%% Parameter search
function f = fun_objective(par,dT,dX,varA,nA,varD,nD)
mustep=par(1);
varstep=par(2);
```

```
for i=1:25
    varterm=dT(i)*varstep+varA(i)/nA(i)+varD(i)/nD(i);
    Li(i)=-log(2*pi)/2-log(varterm)/2-(dX(i)-dT(i)*mustep)^2/(2*varterm);
end

L=sum(Li);

f=-L
end
```
